# Shaping Kale Morphology and Physiology Using Precision LED Light Recipes

**DOI:** 10.1101/2024.10.10.617428

**Authors:** Sabine Scandola, Lauren E. Grubb, Brigo Castillo, Lexyn Iliscupidez, Curtis Kennedy, Nicholas Boyce, Mohana Talasila, R. Glen Uhrig

## Abstract

Light serves as a fundamental factor in plant development, both as an energy source and as an environmental cue. With the advent of light-emitting diode (LED) technology, light can be precisely manipulated to influence key plant traits. Here, we assess effects of light intensity and spectral composition on the growth and physiology of Kale (*Brassica oleracea*). Kale is known for its phenotypic plasticity and nutritional composition, making it a crop well-suited for indoor cultivation either as microgreens or as large leafy plants. Here, we employ a combination of advanced phenotyping, computer vision, gas chromatography-mass spectrometry (GC-MS) metabolomics, and liquid chromatography-mass spectrometry (LC-MS)-based quantitative proteomics to characterize the molecular changes that underpin light-dictated differences in the growth and metabolism of two different commercially grown kale cultivars under different light intensities and spectral compositions. We identify time-of-day and cultivar-specific light intensity and spectral composition-induced changes related to growth, shade avoidance, photosynthesis and several aspects of nutritional composition, including amino acids, glucosinolates and carotenoids. Our results offer a key resource to the plant community and demonstrate the translational potential of light manipulation in tailoring kale growth and nutritional content for enhanced crop productivity and/or health benefits, while simultaneously offering a more cost-effective solution for contemporary agricultural challenges.

**Significance Statement:** The effects of light intensity and spectral composition differentially affect the diel molecular responses of Kale (*Brassica oleracea*). Our results demonstrate the translational potential of light manipulation in tailoring plant growth and nutritional content for enhanced crop productivity.

## Introduction

Kale (*Brassica oleracea*), also known as leaf cabbage, belongs to the large *Brassicaceae* family, which includes multiple agriculturally important crops such as cauliflower, broccoli, and collard greens. Established as a so-called ‘superfood’, this leafy green is rich in fiber, essential vitamins, and antioxidants such as flavonoids, each of which have been shown to positively affect cardiovascular and gastrointestinal health^1^. To date, the health benefits of kale have been heavily studied, with researchers continuing to identify new anti-cancer (e.g., sinigrin, spirostanol)^2^ and neuroprotective (e.g., sulforaphane)^3^ compounds, which neutralize reactive oxygen species. Further, kale demonstrates excellent temperature resilience resulting from its waxy coating and high production of polyamines^4^, rendering it a valuable agricultural crop in the context of climate change^5^.

Kale is also well-suited for indoor and vertical farming systems. Grown either as a microgreen or a fully grown leafy green, kale is a favored choice for horticultural growers as it has a quick harvest cycle with minimal space and resource requirements^6^, offering opportunities for fresh, locally sourced produce^7^. Indoor / vertical farms have strongly co-evolved with new technologies, with the promise of fresh and locally grown food largely relying on reduced production costs through technological advancements^8^. In this context, LED technology has revolutionized indoor / vertical farming, providing a wide range of energy-efficient and versatile lighting options. For example, LED lighting systems offer new abilities to fine-tune the spectral composition and intensity of applied light relative to conventional lighting systems containing either incandescent, fluorescent, metal halide or high-pressure sodium bulbs^9^.

Light intensity and spectral composition are critical factors for plants. Light intensity is important for photosynthesis, which drives plant metabolism^10^, whereas spectral composition, detected by different plant photoreceptors such as phytochromes and cryptochromes, fine-tunes processes such as plant growth (e.g., biomass) and development (e.g., flowering)^11^. However, despite our understanding of kale at a phenotypic level, we lack knowledge of how light impacts kale at the molecular level, limiting our ability to develop enhanced horticultural practices and / or genetics through targeted breeding or biotechnological approaches.

To date, elementary changes in spectral composition have been shown to affect kale phenotypic and metabolic profiles^12^. Blue light enhances growth and nutritional qualities^13^, while red light affects kale pre- and post-harvest by altering nutrient and chlorophyll content^14^. Similar to red light, far-red light enhances photosynthetic rates through the Emerson effect^15^. Beyond the application of specific light types, overall spectral composition also plays an essential role in how phytochromes perceive and transduce light signals^16^, offering opportunities to build precise light recipes that maximize traits of interest. Use of precision LED technologies now offers a new ability to fully explore this possibility.

To explore how we could modulate the growth of commercially available kale genetics, with a particular interest for use in vertical farming systems which typically employ lower light intensities, we endeavored to precisely resolve the impacts of light intensity and spectral composition on kale growth, development and nutritional content by contrasting the diel molecular changes in two related kale cultivars: Dwarf curled scotch (K3) and Red Scarlet (K9). While there are many commercially available kale cultivars, each with unique morphologies, growth parameters and nutritional characteristics^17^, our previous study examining nine of these kale cultivars found that K3 and K9, both *Brassica oleracea var. sabellica*, showed the most diversity in their phenotypes and molecular characteristics^17^, allowing the results presented here to capture how the application of light can impact kale at the phenotypic-, proteomic- and metabolomic-level. As both the proteome and metabolome are known to undergo substantial diel changes^17^ and indoor / vertical farming offers opportunities for precise time-of-day harvesting, we took a diel experimental approach to analyze the molecular effects of light intensity and spectral composition by harvesting tissues for analysis at end-of-day (ED, ZT11 (zeitgeber)) and end-of-night (EN, ZT23) time-points. Understanding how external factors such as time-of-day intersect with differences in light intensity and quality to impact kale growth and underlying molecular mechanisms is essential for determining optimal indoor / vertical farming growth conditions and harvesting times to extract and maximize nutrition and value, as well as inform efforts in breeding more productive and nutritious kale varieties.

## Results and Discussion

### Light intensity and spectral composition influence kale morphology

We first investigated the effects of light intensity and spectral composition on the morphology of kale. For our light intensity experiments, plants were grown under three different light intensities: 75, 125 and 175 PPFD (**Figure 1A**). These light intensities were chosen based on conditions of a typical vertical farming operation, which generally employ a PPFD of 150-250^18^. Top-view cameras were used to take images every 5 min, and these were averaged over a 1 h period between ZT6 and ZT7 to minimize noise at 22 dpi. This revealed a relatively linear relationship between light intensity with plant fresh weight and surface area in both cultivars at 35 days post-imbibition (dpi) (**Figure 1B-C**). Consistent with our observations, previous studies have reported a linear relationship between light intensity and fresh weight in pea microgreens^19^, as well as kale, cabbage, arugula and mustard^20^. Some of these studies used PPFD up to 600 µmol·m^-2^·s^-1^, with increases in fresh weight occurring even at the upper PPFD values, however, light intensities in vertical farming operations are typically around 150-200 PPFD^21–23^, suggesting our highest light intensity of 175 PPFD is aligned with industry standards. It is important to note for our further analyses that the plants grown at different light intensities were at different developmental stages, as those grown under the higher PPFD have more leaves and are more advanced in development. Next, we examined the effect of light intensity on diel leaf anthocyanin content, which provides plant coloration to attract pollinators, while also functioning as an antioxidant with benefits to human health^24^. Although anthocyanins could not be detected in K3, the K9 cultivar, which is known to produce anthocyanins^25^ typically showing red colouration at later stages of development, did not show a significant difference in anthocyanin concentration with increasing light intensity at the PPFDs tested (**Figure 1E**). Anthocyanins are typically produced in response to stressful conditions^26^; thus, it is possible that our use of lower light intensities than that experienced outdoors, combined with the stage of growth, resulted in lower anthocyanin content than might have been expected.

**Figure 1.**
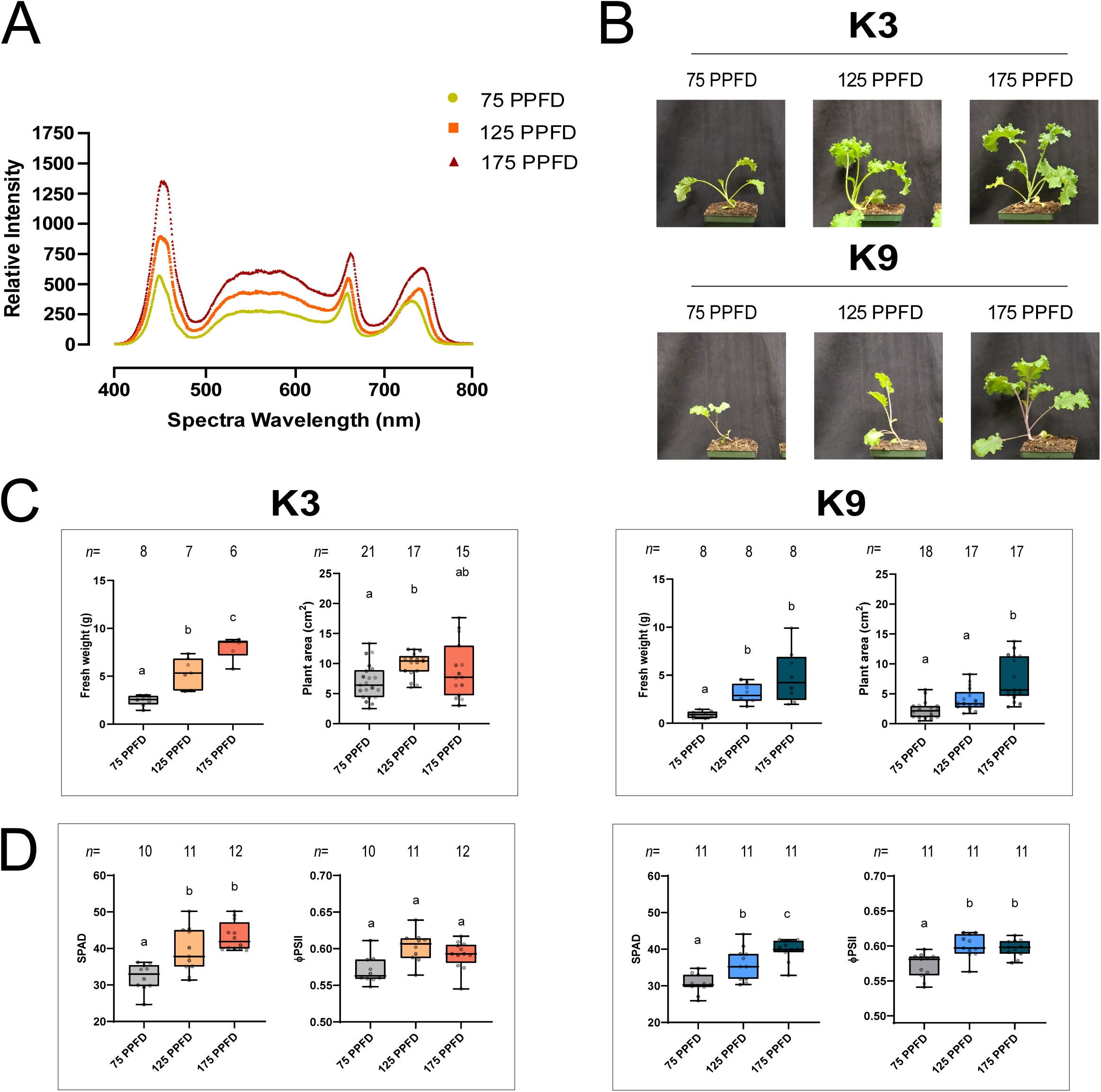
**Effect of light intensities on kale growth and development.** A. Light spectra of different light intensity (75, 125 and 175 PPFD). B. Pictures of K3 and K9 plants at 35 dpi grown under different light intensities (75, 125, and 175 PPFD). C. K3 and K9 fresh weight at 35 dpi (n≥6), leaf area at 22 dpi (n≥15). D. Relative chlorophyll content (SPAD) and photosynthetic parameters (Phi2) measured with MultispeQ at 35 dpi (n≥10). Letters shows significant differences using a one-way ANOVA and Tukey’s post-hoc test (adjusted p-value <0.05).

Our spectral composition experiments consisted of modulating the ratio of red to blue (R/B) and red to far-red (R/FR) light, while maintaining an otherwise balanced spectra and a light intensity of 125 PPFD (**Figure 2A**). Previous studies have shown a high R/B ratio promotes plant growth^27^, whereas a low R/B ratio promotes accumulation of beneficial metabolites^28^. Here we found that increasing the R/B light ratio increased fresh weight in K3 with no corresponding change in surface area, whereas K9 exhibited no change in fresh weight, but an increase in plant surface area with increasing R/B and R/FR ratios (**Figure 2B-C**), indicating that red light may be particularly important for K9 growth. Supporting this possibility are observed increases in dry mass and seedling area specifically in red kale varieties under elevated red light^29^, suggesting spectral composition impacts on growth can depend on the cultivar. This emphasizes the need to generate cultivar-specific molecular response data, such as that investigated here, to uncover unique breeding opportunities and inform growers choosing genetics for indoor / vertical farming. We additionally observed changes in K9 anthocyanin content with modulated spectral composition (**Figure 2E**). Consistent with previous reports, an increase in R/B induced a decrease in anthocyanin content, whereas an increase in R/FR stimulated anthocyanin production at both time-points (**Figure 2E**). This indicates that changing light intensity and spectral composition impacts both kale morphological characteristics and anthocyanin levels.

**Figure 2.**
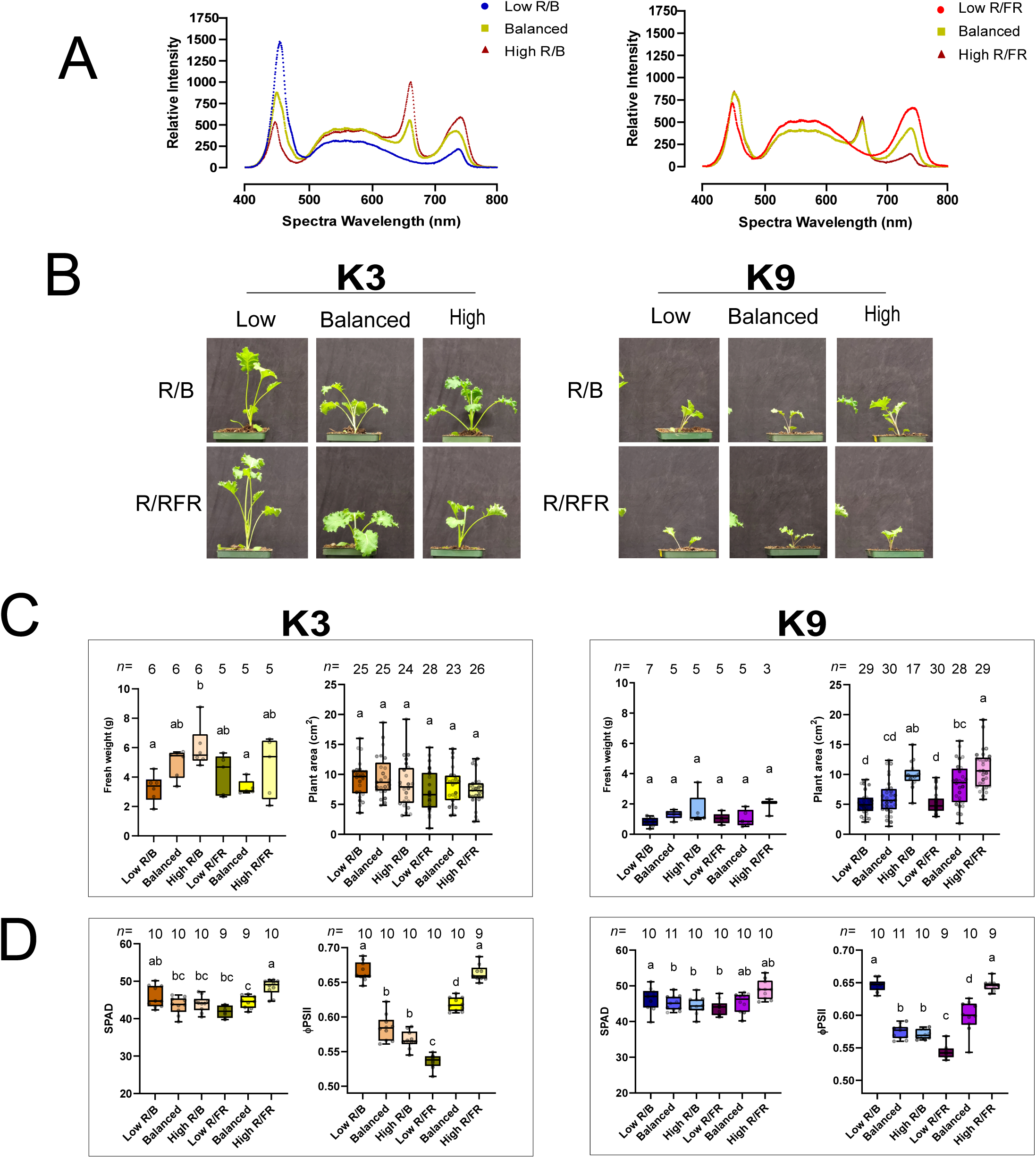
**Effect of light spectra on kale growth and development.** A. Light spectra of the different light ratios (low R/B, balanced and high R/FR). Light intensity is 125 PPFD for all spectra treatment. B. Pictures of K3 and K9 plants at 35 dpi grown under different light spectra (low, balanced, high R/B and R/FR). C. K3 and K9 weight at 35 dpi (n≥3), leaf area at 22 dpi (n≥17) D. Relative chlorophyll content (SPAD) and photosynthetic parameters (Phi2) measured with MultispeQ at 35 dpi (n≥9) at different light spectra (low, balanced, high R/B and R/FR). Letters shows significant differences using a one-way ANOVA and Tukey’s post-hoc test (adjusted p-value <0.05).

### Modulating light intensity and spectral composition impacts kale photosynthetic performance

We next measured light intensity- and spectra-induced changes in a suite of photosynthetic parameters alongside relative chlorophyll levels using a MultispeQ device^30^ (detailed in **Table 1**). Here we noted that ΦPSII, which represents how much light energy is absorbed by photosystem II^30^, and chlorophyll content (SPAD), were both significantly impacted by changes in light intensity and spectral composition. Increases in light intensity (**Figure 1D**) and R/FR (**Figure 2D**) increased SPAD in both cultivars, whereas increasing R/B decreased SPAD in both cultivars (**Figure 2D**). Similarly, we find elevated ΦPSII under increasing light intensity, but in the K9 cultivar only (**Figure 1D**), while elevated ΦPSII was found in both cultivars with increasing R/FR (**Figure 2D**). Conversely, an increasing R/B ratio resulted in decreased ΦPSII across both cultivars (**Figure 2D**). Our findings of elevated ΦPSII with increasing light intensity aligns with previous observations in lettuce where the highest ΦPSII was found at 200 PPFD across a range of 100 to 800 PPFD, which is comparable to our 175 PPFD treatment^31^. Furthermore, blue light is found to be more important than red light in modulating photosystem I and II activity and photosynthetic electron transport capacity in cucumber^32^, consistent with our increasing R/B ratio showing lower SPAD and ΦPSII. Overall, our results indicate that kale photosynthetic parameters and chlorophyll content can be modulated by changes in light intensity and spectral composition.

**Table 1.**
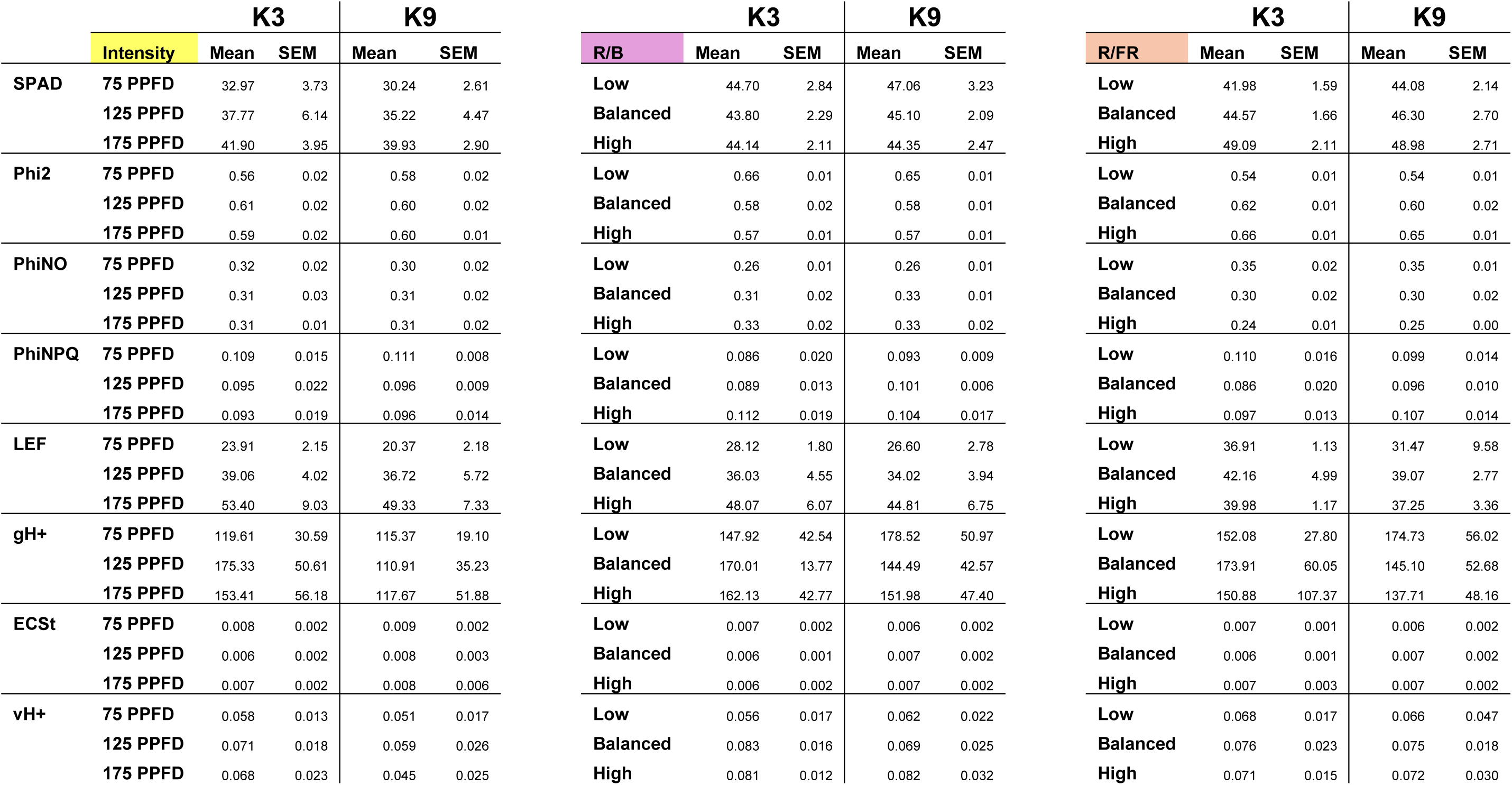
Effect of light intensity and spectra on photosynthetic parameters measured with the MultispeQ (PhotosynQ Inc.). Relative chlorophyll content (SPAD) and photosynthetic parameters (Phi2, PhiNO, PhiNPQ, LEF, gH+, ECSt, vH+) were measured with the MultispeQ at 35 DPI (n=10).

### Analysis of total proteome changes with altered light intensity and spectral composition

To contextualize the observed phenotypic changes, we used quantitative proteomics to assess the protein-level changes in both K3 and K9 across the same light intensities and spectral compositions. This analysis quantified a total of 5,493 proteins across our intensity experiments, and 6,527 proteins across spectral composition experiments (**Supp Table 1**). We queried this data for significantly changing proteins by comparing each light intensity or spectral composition to the lowest treatment condition. Thus, 125 PPFD and 175 PPFD were compared to 75 PPFD (**Figure 3A; Supp Table 1**) and ‘balanced’ and ‘high’ spectral compositions were compared to the ‘low’ spectral composition (**Figure 3B-C; Supp Table 1**).

**Figure 3.**
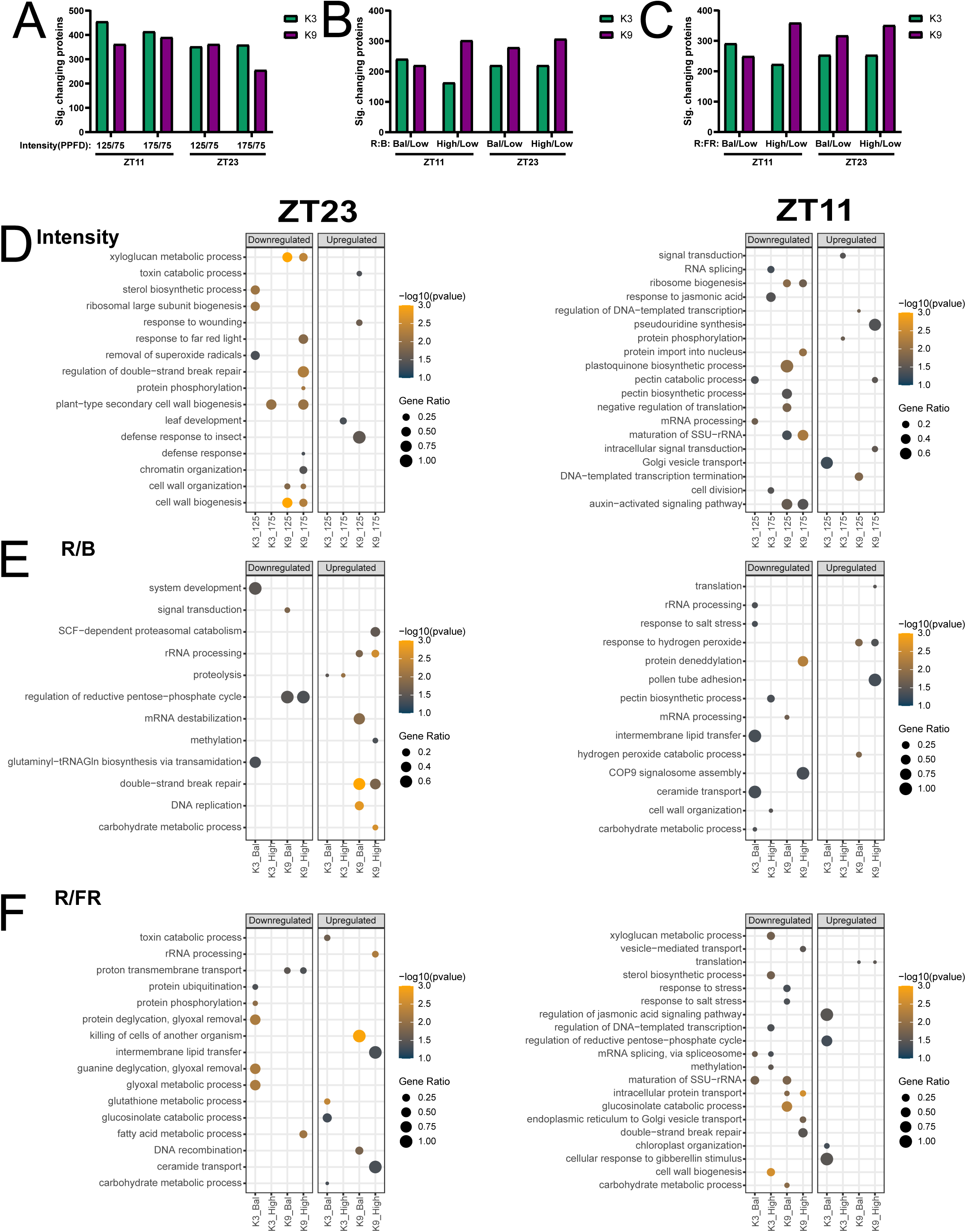
**Analysis of significantly changing proteins with changing light intensity and spectral composition.** A-C. Number of proteins with a significant change in abundance with increase in light intensity from 75 PPFD (A), from low R/B (B) and from low R/FR (C) determined based on Bonferroni-corrected p-value <0.05 from 4 replicates. D-F. Dot plot representation of enriched GO terms for biological process of significantly changing proteins (q-value < 0.05; Log2FC > 1 or < −1) that are up- or down-regulated with increasing light intensity (D), increasing R/B ratio (E), or increasing R/FR ratio (F) at ZT23 (left) or ZT11 (right). Dot size indicates the gene ratio (number of proteins in the changing in the condition/total quantified proteins in the category in the study). Dot colour represents the −Log10 (p-value). Only GO terms with p-value <0.05 and a total count < 500 are included.

Using these significantly changing proteins (Log2FC < −1 or > 1), we undertook gene ontology (GO) enrichment analysis to resolve biological processes underpinning plant response(s) to each changing light intensity and spectral composition (**Figure 3D-F**). This analysis revealed an enrichment in biological processes related to cell wall and cell cycle, response to oxidative stress and hormone signaling (auxin, gibberellin, jasmonic acid), along with extensive changes in multiple aspects of metabolism, including fatty acid metabolism, carbohydrate metabolism, and glucosinolate production. We next refined our GO analysis results by performing an association network analysis using STRING-DB (https://string-db.org/; edge score > 0.9 for Intensity and R/FR and > 0.8 for R/B) to define hubs of associated proteins exhibiting cultivar or time-of-day responses. To do this, we used the significantly changing proteins (Log2FC <-1 or >1) induced by a change in light intensity (**Figure 4**), R/B (**Figure 5**) and R/FR (**Figure 6**). As a closely related kale relative, *Arabidopsis thaliana* (Arabidopsis), has more extensive bioinformatic resources available. Therefore, we first converted the *B. oleracea* gene identifiers for all quantified proteins in the study to Arabidopsis gene identifiers (AGI) using a combination of UniProt (https://www.uniprot.org/) and Ensembl Biomart (https://plants.ensembl.org/biomart). This identified unique AGIs for 90% of the significantly changing proteins in our intensity experiments (2395/2662), along with 88.9% (2222/2499) and 89.4% (2520/2819) of proteins in our R/B and R/FR spectra experiments; respectively. Along with validating enriched biological processes, the association network further revealed changes in amino acid metabolism, phenylpropanoid metabolism, photosynthesis and pigment metabolism, proteolysis and RNA splicing.

**Figure 4.**
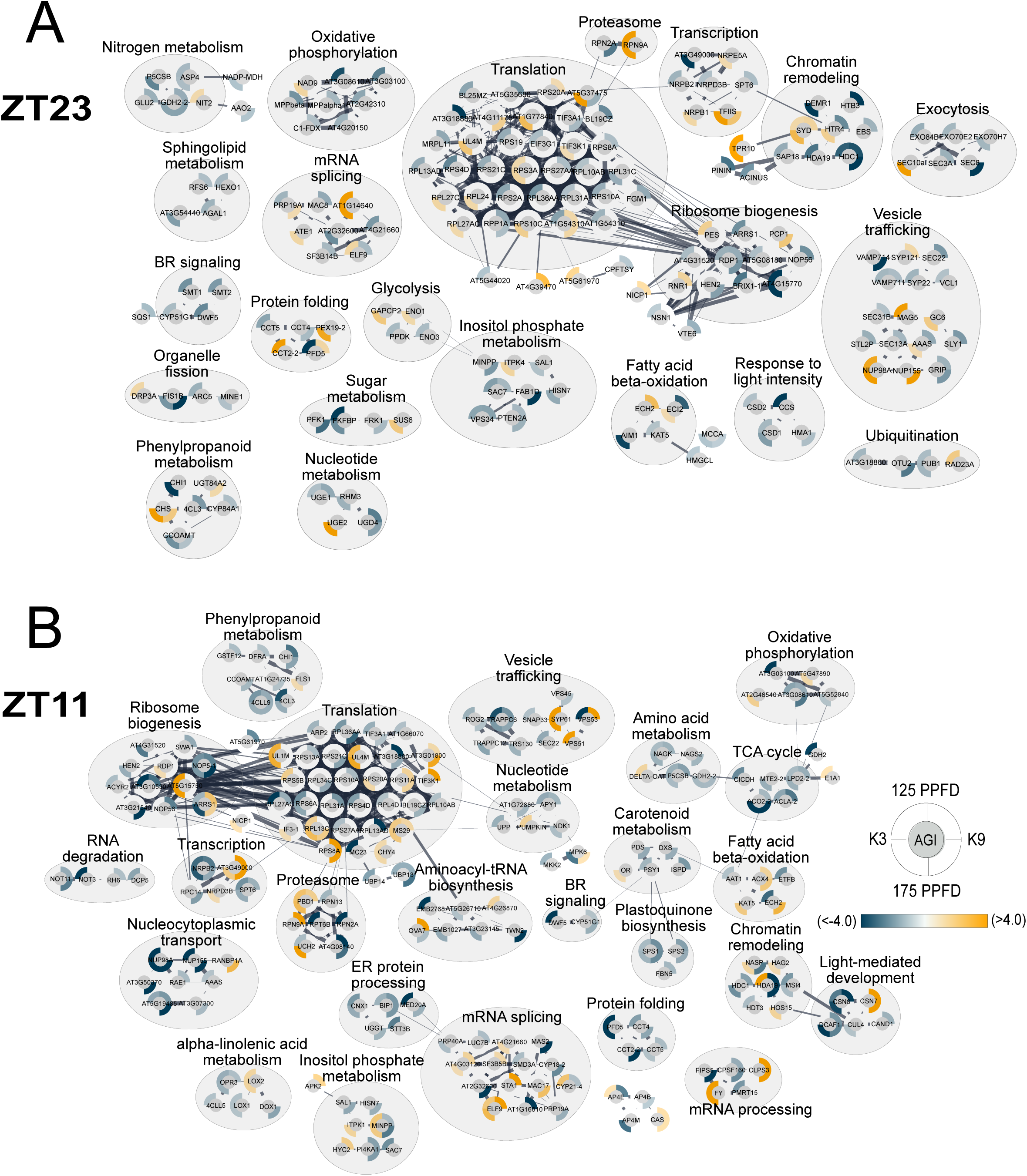
**Association network analysis of significantly changing proteins with increasing light intensity.** An association network analysis was performed using STRING-DB to depict significantly changing proteins (q-value < 0.05; Log2FC > 1 or < −1) in K3 and K9 kale under increasing light intensity at ZT23 (A) and ZT11 (B). Edge thickness indicates strength of connection between the nodes. Minimum edge threshold was set to 0.9. Protein nodes are labelled either by primary gene name or Arabidopsis gene identifier (AGI). Outer circle surrounding each node represents the standardized relative Log2FC of the indicated significantly changing protein with an increase from 75 PPFD to either 125 PPFD or 175 PPFD in K3 or K9 as indicated by the legend. The scale of blue to yellow indicates the relative decrease or increase in abundance, respectively from 75 PPFD to either 125 PPFD or 175 PPFD. Node groupings are indicated by a grey circle representing proteins involved in the same biological process.

**Figure 5.**
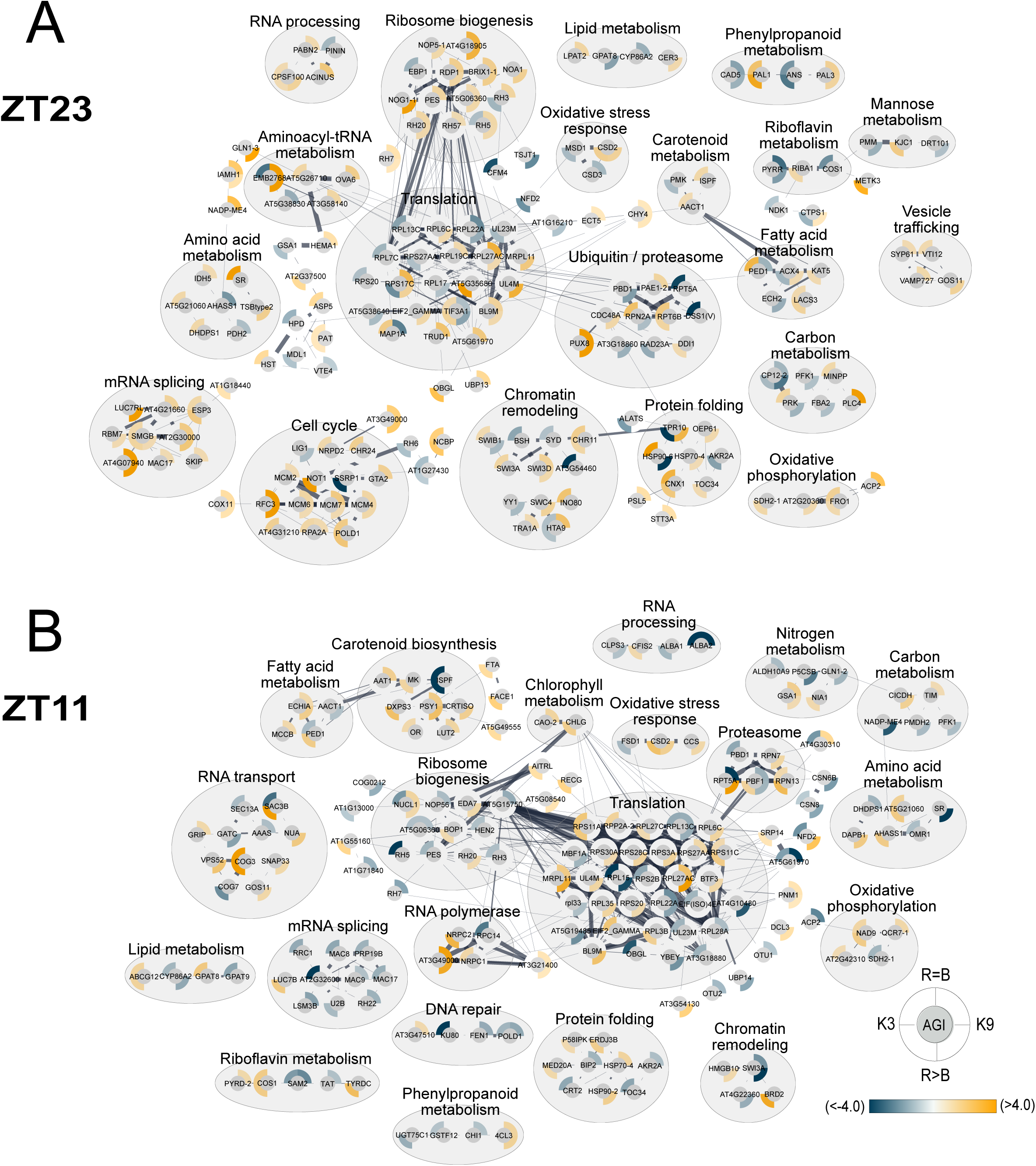
**Association network analysis of significantly changing proteins with increasing R/B light spectra.** An association network analysis was performed using STRING-DB to depict significantly changing proteins (*q-value* < 0.05; Log2FC > 1 or < −1) in K3 and K9 kale under increasing R/B spectral ratio at ZT23 (A) and ZT11 (B). Edge thickness indicates strength of connection between the nodes. Minimum edge threshold was set to 0.8. Protein nodes are labelled either by primary gene name or Arabidopsis gene identifier (AGI). Outer circle surrounding each node represents the standardized relative Log2FC of the indicated significantly changing protein with an increase from R<B to either R=B or R>B in K3 or K9 as indicated by the legend. The scale of blue to yellow indicates the relative decrease or increase in abundance, respectively from R<B to either R=B or R>B. Node groupings are indicated by a grey circle representing proteins involved in the same biological process.

**Figure 6.**
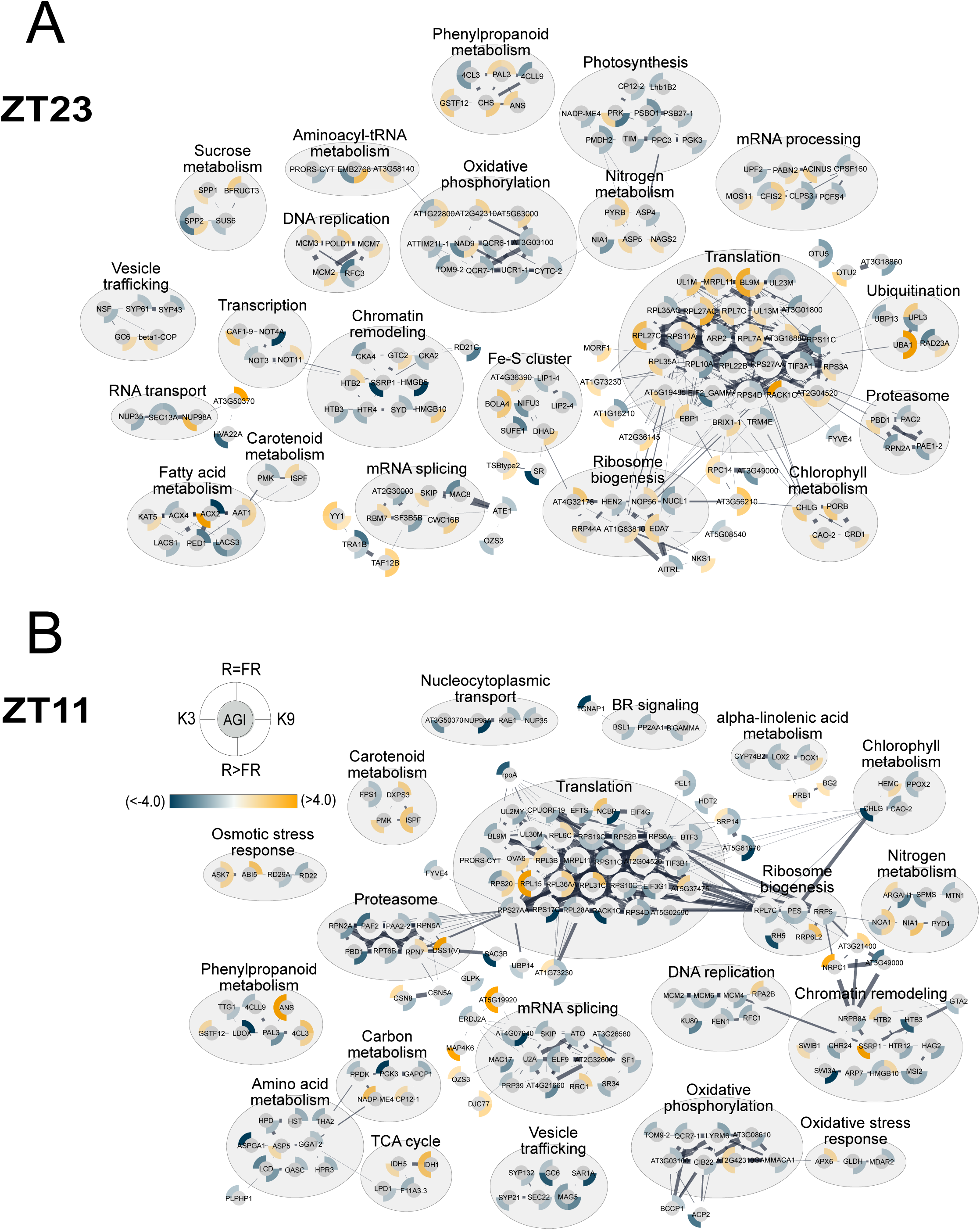
**Association network analysis of significantly changing proteins with increasing R/FR light spectra.** An association network analysis was performed using STRING-DB to depict significantly changing proteins (q-value < 0.05; Log2FC > 1 or < −1) in K3 and K9 kale under increasing R/FR spectral ratio at ZT23 (A) and ZT11 (B). Edge thickness indicates strength of connection between the nodes. Minimum edge threshold was set to 0.9. Protein nodes are labelled either by primary gene name or Arabidopsis gene identifier (AGI). Outer circle surrounding each node represents the standardized relative Log2FC of the indicated significantly changing protein with an increase from R<FR to either R=FR or R>FR in K3 or K9 as indicated by the legend. The scale of blue to yellow indicates the relative decrease or increase in abundance, respectively from R<FR to either R=FR or R>FR. Node groupings are indicated by a grey circle representing proteins involved in the same biological process.

Interestingly, several of the enriched biological processes could be attributed to the plant shade avoidance response, or relief thereof, including the changes in hormone signaling such as brassinosteroid (BR) and auxin signaling, cell wall organization / biogenesis and photosynthesis. Plant shade avoidance response is associated with conditions that mimic shading, including low light intensity, low blue light, and low R/FR ratios^33^ and has been linked to increased BR^34–36^ and auxin signaling^37^; both of particular importance for shade-induced hypocotyl elongation. Within our dataset, we find a down-regulation of BR signaling in both cultivars with increasing light intensity and R/FR at ZT11 (Figure 3, 4, 6), and a down-regulation of auxin-activated signaling pathway with increasing intensity at ZT11 (Figure 3). Both observations are consistent with a relief of shade avoidance responses with our increasing intensity experiments, and an induction upon growth under decreased blue light.

Further, BR shade avoidance response is related to changes in cell wall structure, particularly through the action of XYLOGLUCAN ENDOTRANSGLUCOSYLASE / HYDROLASES (XTHs) and EXPANSINS (EXPAs) proteins, with cell wall modification promoting elongation of the hypocotyl to drive growth toward light^38^. Our data find a multitude of cell wall proteins changing in abundance in response to both light intensity and spectral composition, potentially driving shade relief-like responses in both kale cultivars. We specifically note a down-regulation of cell wall organization proteins (Figure 3), primarily XTHs, EXPAs, and PECTINESTERASES (PMEs) with increasing light intensity and R/FR ratio, combined with an up-regulation of XTHs with increasing R/B, which is consistent with changes in the plant perception of shade and hypocotyl elongation. Interestingly, more members of the XTH family were downregulated in K9 with increasing light intensity and K3 with increasing R/FR, suggesting K9 may be more sensitive to changes in intensity and K3 more sensitive to changes in red light wavelengths, a finding that agrees with our growth and morphology observations.

### Analysis of metabolite changes with modified light intensity and spectral composition

With a multitude of metabolic pathways enriched in our GO and STRING-DB analyses, we next used the Arabidopsis orthologs of the kale proteins exhibiting significant changes in abundance as a result of increasing intensity, R/B and R/FR, to perform metabolic pathway analyses using the Plant Metabolic Network (PMN; https://plantcyc.org; **Supp Tables 2-4**). Here, we find enriched metabolic pathways that further corroborated our GO and STRING-DB association network analyses, resolving pathways for photosynthesis and respiration, amino acid metabolism, pigment metabolism, glycolysis, carbohydrate metabolism, oxidative stress responses, glucosinolate production and sterol biosynthesis. We next sought to determine how these changes in metabolic pathway protein abundance translate into metabolite compositional changes. To do this we used GC-MS to quantify metabolite changes in each kale cultivar under each light intensity (**Figure 6; Supp Figure 3; Supp Table 5**), R/B ratio (**Figure 7A; Supp Figure 4; Supp Table 6)** and R/FR ratio (**Figure 7B; Supp Figure 5; Supp Table 6)**. Through this analysis we were able to quantify a range of amino acids (16), fatty acids (3), organic acids (11), polyamines (2), sterols (2), sugars (5), and sugar alcohols (2) across all samples for comparative analysis.

**Figure 7:**
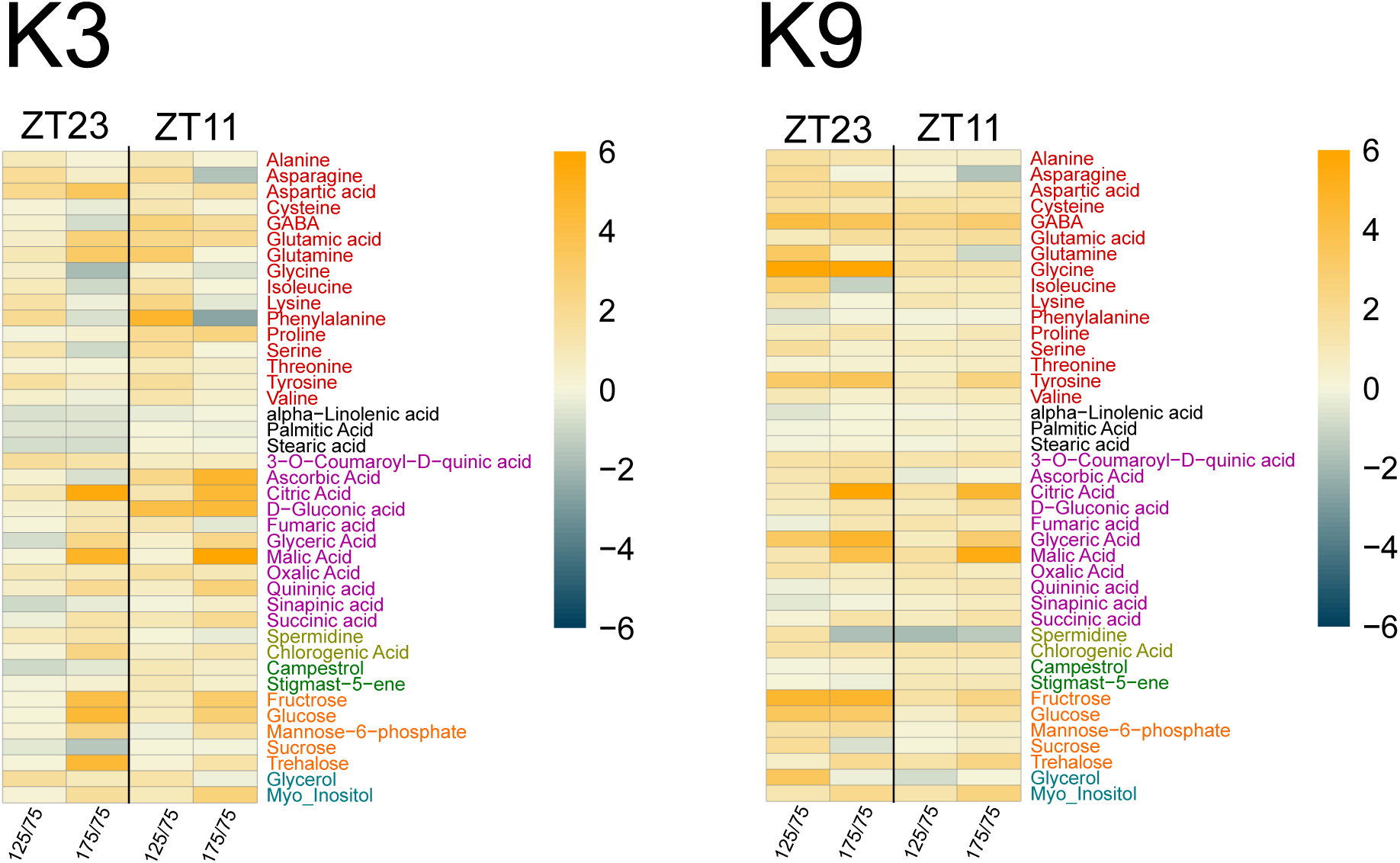
**GC-MS analysis of kale cultivars K3 and K9 under different light intensities (75 PPFD, 125 PPFD and 175 PPFD) and at different time of the day (ZT11 and ZT23).** Heatmap of relative cultivars metabolite changes at ZT11 and ZT23 for 125 PPFD/75 PPFD and for 175 PPFD/75PPFD. Scale represents log2FC, n=4

*Amino acid metabolism* - Quantitative proteome analysis identified a significant change in amino acid metabolic proteins in both kale cultivars in response to changing light intensity and spectral composition (**Figures 7-8; Supp Figures 3-5; Supp Tables 5-6**). Correspondingly, our metabolite analysis found extensive changes in the glutamate-derived family of amino acids, including glutamic acid, glutamine, GABA, proline and arginine. These amino acids typically account for the highest proportion of amino acids in plants, with glutamic acid and proline present at the highest quantity in kale^39^. Glutamate metabolism is also a critical link between nitrogen and carbon metabolism through the TCA cycle, with several products of glutamate catabolism, including proline and GABA, associated with plant stress response^40,41^.

**Figure 8:**
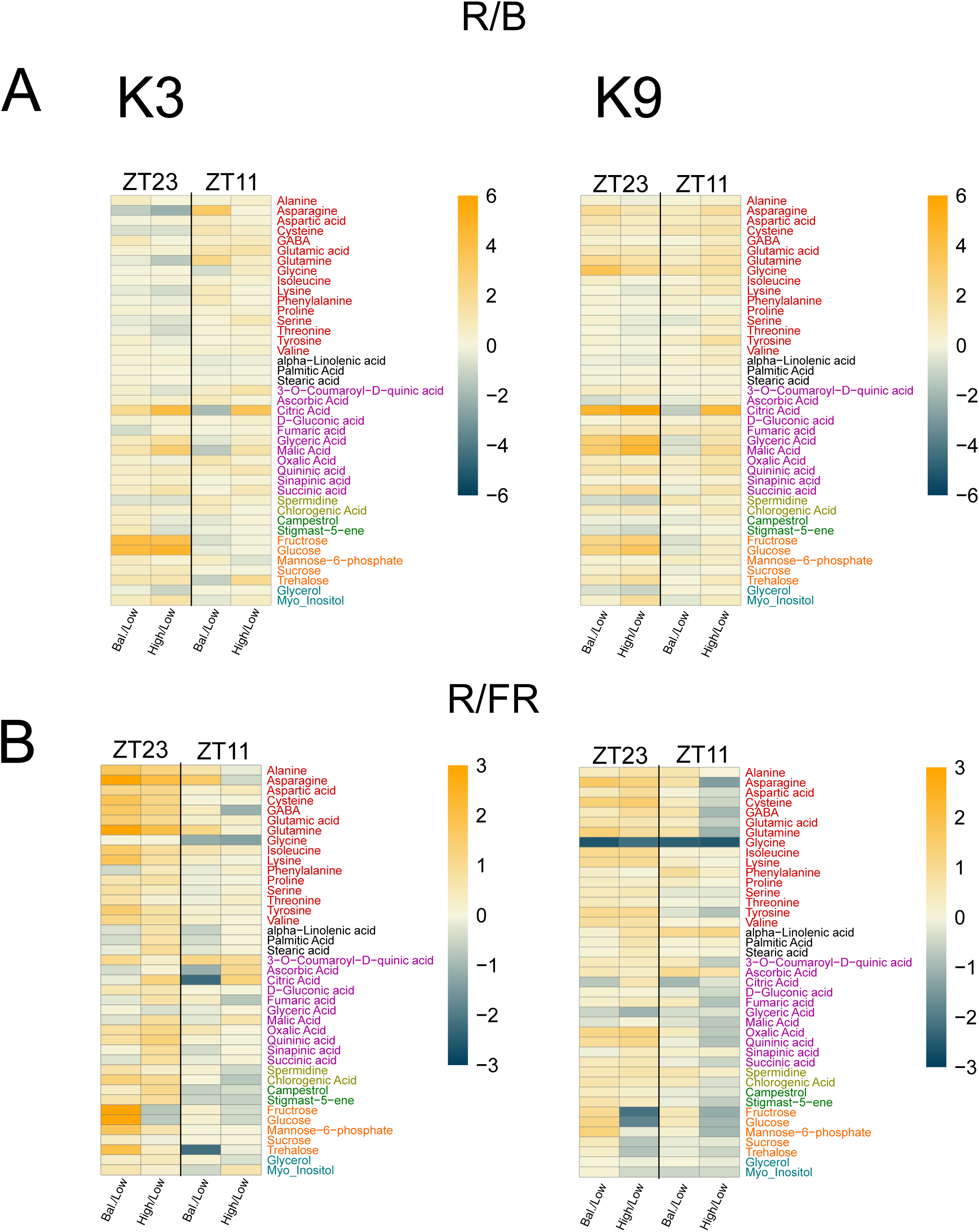
**GC-MS analysis of kale cultivars K3 and K9 under different spectral compositions.** A) Heatmap of relative cultivars metabolite changes at ZT11 and ZT23 for changing R/B ratio Balanced/Low and High/Low. Scale represents log2FC, n=4 B) Heatmap of relative cultivars metabolite changes at ZT11 and ZT23 for changing R/FR ratio Balanced/Low and High/Low. Scale represents log2FC, n=4

**Figure 9.**
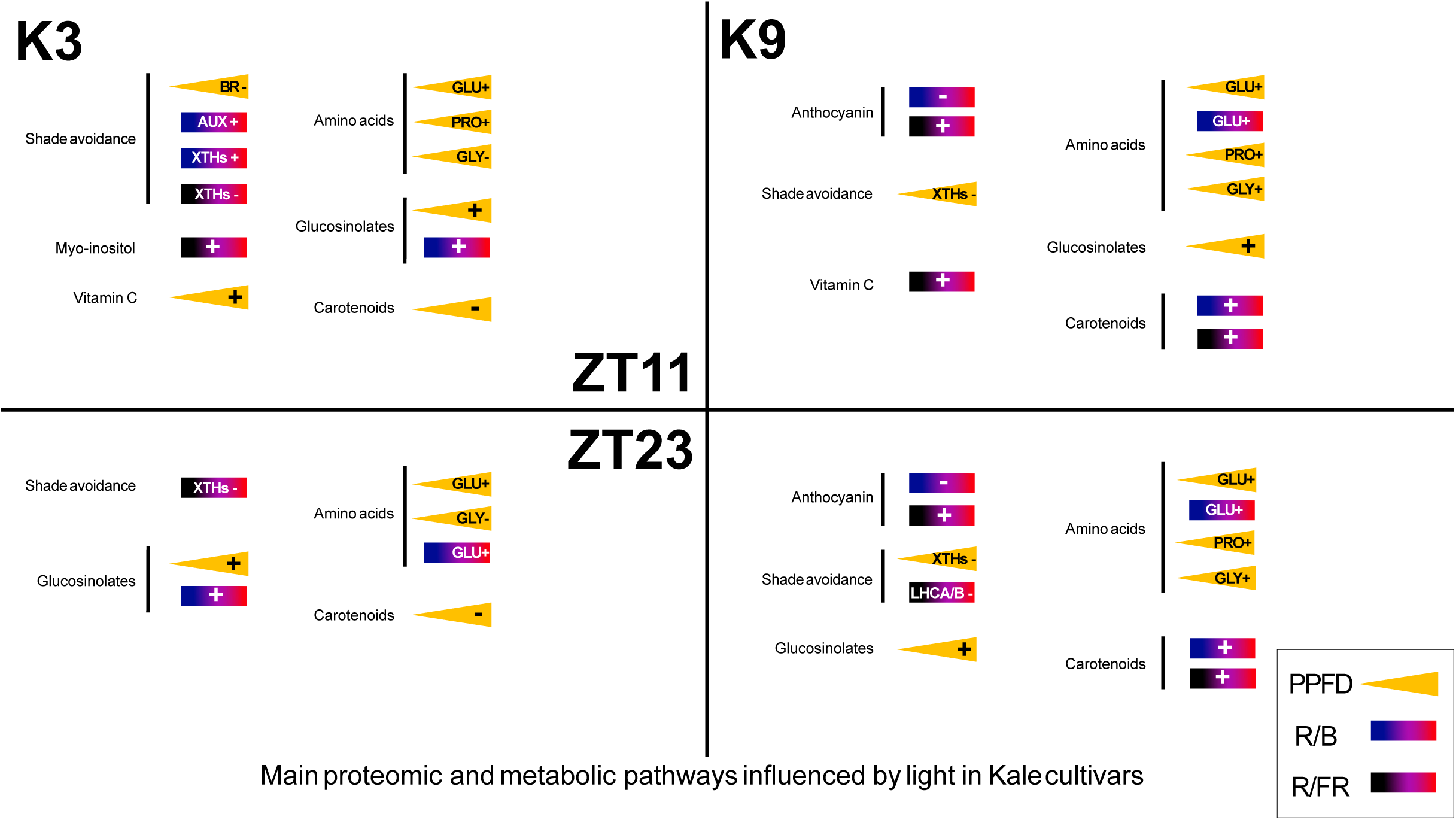
**Summary of proteomic and metabolic changes with varying light intensity and spectra.** Summary of main proteomic and metabolic pathways influenced by light in kale cultivars K3 and K9 at ZT11 and ZT23 with increasing PPFD, R/B and R/FR.

In both cultivars, we observed a general increase in glutamic acid and proline at both time-points with increasing light intensity. GABA increased at ZT11 in both cultivars, followed by a decrease in GABA levels at ZT23 in K3 and an increase in K9. Increased glutamic acid abundance with increasing light intensity coincided with our proteome-based metabolic pathway analysis, where we identified down-regulation of proteins involved in glutamate catabolism (Figure 4). This included reduced abundance of GLUTAMATE DEHYDROGENASE1 (GDH1; A0A0D3EEJ0) and GDH2 (A0A0D3AI31), which convert glutamate to glutamine, DELTA-1-PYRROLINE-5-CARBOXYLATE SYNTHASE B (P5CSB; A0A0D3BXP3), which is involved in proline biosynthesis, N-ACETYLGLUTAMATE SYNTHASE (NAGS; A0A0D3A172) and N-ACETYLGLUTAMATE KINASE (NAGK; A0A0D3BX89), which are both involved in ornithine biosynthesis, and CARBAMOYL-PHOSPHATE SYNTHASE ALPHA (CARA; A0A0D3CZA9), which drives arginine biosynthesis. We find this trend in both K3 and K9 and at both time-points (Figure 4), suggesting increasing the light intensity in our experiments may relieve stress responses associated with low light. The increase in proline concentration under higher light intensities (Figure 7; Supp Figure 3), however, may suggest sufficient levels of proline, with the down-regulation of P5CSB preventing further proline accumulation.

Similarly, our increased R/B ratio data revealed changes associated with proline biosynthesis and glutamate metabolism both K3 and K9, while glutamic acid levels in each kale cultivar increased at both time-points. Proline levels were not responsive to increasing R/B, however, glutamine peaked under balanced light condition at ZT11 and decreased at ZT23. Alternatively, increasing R/FR induced large changes in the metabolite content of glutamate-family amino acids. Interestingly, we find peak glutamic acid, glutamine and GABA levels in both K3 and K9 under balanced light conditions. This may indicate that balanced light conditions are associated with the most balanced C/N metabolism during the day, but highlight how changing spectral composition can have wide-ranging metabolite effects.

Aspartate-derived amino acids are also of great importance for human nutrition as they cannot be synthesized by the human body and must be obtained through diet^42^. This includes essential amino acids lysine, threonine, methionine and isoleucine. Of particular interest is lysine, as it is generally of very low abundance in plants^43^. Within our proteome metabolic pathway analysis, lysine biosynthesis is enriched among down-regulated proteins as light intensity increases (**Supp Table 2**), particularly in K3 at ZT11, with significantly reduced abundance of ASPARTATE KINASE (AK2) and of DIHYDRODIPICOLINATE SYNTHASE1 (DHDPS1); both involved in lysine biosynthesis. However, within our metabolite data (**Figure 7; Supp Figure 3**), there is a significantly higher amount of lysine present in K3 under 125 PPFD than either 75 or 175 PPFD, suggesting more complex regulation. Interestingly, regulatory complexity in the production of lysine has been previously observed by biotechnology efforts to increase plant lysine production^44^. Here, it was found that lysine allosterically inhibits DHDPS1 activity^44^ as well as impact the abundance of other aspartate-derived amino acids, such as threonine^45^. Our metabolite data suggests that lysine levels can be modulated by changing light intensity and spectra, with peak lysine levels found at 125 PPFD for both cultivars at ZT23 (**Figure 7; Supp Figure 3**), and a peak lysine levels under balanced spectral conditions in K3 at ZT23 (**Figure 8; Supp Figure 4-5**). Thus, our results suggest the difficulty in engineering lysine levels in kale may be overcome through careful modulation of light intensity and spectral composition.

Finally, we also noted cultivar-specific differences in glycine content with increasing intensity (**Figure 7; Supp Figure 3**). We observed decreased glycine abundance with increasing light intensity in K3, but increasing glycine levels in K9. Glycine uptake has previously been reported to increase with increasing light intensity in pakchoi^46^. Previously, we observed differential accumulation of glycine to represent a major difference between kale sub-groups, including K3 and K9^17^, which could correspond to differences in nitrogen metabolism strategies. Although we did not observe much change in serine content, through which glycine is derived, both glycine and serine are key to the functioning of primary metabolism and photorespiration^47^, and thus this difference may point to cultivar-specific mediation of primary metabolism responses to differing light intensity.

*Carbohydrate metabolism* – Our proteomic analysis revealed extensive changes in carbohydrate metabolic proteins upon changes in both light intensity and spectral composition (**Figures 4-5**). We also find a differential abundance of several sugars in our metabolite analysis, including fructose, glucose, mannose-6-phosphate (M6P) and trehalose. Our proteomic data reveals a general down-regulation of proteins involved in glycolysis and starch degradation, as well as trehalose biosynthetic proteins with increasing light intensity (**Figure 4**). As trehalose has been associated with shade avoidance^48^ and regulation of the circadian clock^49^, this is consistent with the low light intensities associated with shade as we increase the light intensity from 75 to 175 PPFD. We also find that increasing R/B down-regulates proteins associated with glycolysis in both cultivars, in addition to mannose metabolism in the K9 cultivar only (**Figures 3, 5**). This was consistent with the lowest M6P levels found in K9 under balanced light conditions (**Figure 7; Supp Figure 4-5**). Finally, increasing R/FR was associated with the down-regulation of glycolytic proteins in both cultivars, as well as myo-inositol biosynthetic proteins in K9 (**Figure 3, 5**). Conversely, myo-inositol biosynthesis was up-regulated in K3 with increasing R/FR at ZT11 (**Figure 3, 5)**, and was previously reported as a metabolite that differentiates kale genetics^17^. With myo-inositol being involved in a number of key biological processes, including auxin biosynthesis, membrane biogenesis, light and abiotic stress responses^50^, its light-driven modulation likely underpins several of the morphological responses we observe.

*Biosynthesis of other nutritional compounds* - In addition to amino acids and carbohydrates, our metabolite and proteome analyses identified light intensity- and spectral composition-induced changes in several other compounds beneficial for human nutrition, with some exhibiting cultivar and time-of-day specific differences. Specifically, we find an increase in ascorbic acid (vitamin C) with increasing light intensity, particularly in K3 at ZT11 (Figure 7; Supp Figure 3). Vitamin C has antioxidant properties^51^, and this finding corresponds with the enrichment of superoxide metabolic processes amongst up-regulated proteins in K3 at ZT11. In addition, we find that our R/FR experiments reduce vitamin C in K3 under balanced spectral conditions (Figure 8; Supp Figure 4), while increasing in abundance with an increase in R/FR (Figure 8; Supp Figure 5), indicative of the balanced light condition having the lowest oxidative stress. Conversely, we find ascorbic acid increasing in abundance with increasing R/FR in K9 (Figure 8; Supp Figure 5), suggesting different responses to oxidative stress amongst kale cultivars, potentially coinciding with the increased anthocyanin production capabilities of K9. For example, it is possible that K3 deals with oxidative stress by increasing ascorbic acid and K9 by increasing anthocyanin content. The K3 ascorbic acid content is consistent with previous findings in green pepper fruit, where increased red and far-red light both increased ascorbic acid content^52^.

In addition to ascorbic acid, anthocyanins are a natural source of antioxidants, beneficial for human health^53,54^, with the generation of anthocyanins typically driven by light^55^. In our study, we were only able to detect anthocyanins within the K9 cultivar, with no significant difference under the changing light intensity, but a significant decrease with increasing R/B, and a significant increase with increasing R/FR. Within our metabolite analysis, we observe changes associated with anthocyanin and phenylpropanoid metabolism: including chlorogenic acid and quininic acid with changing R/B, and sinapinic acid, chlorogenic acid, quininic acid and 3-coumaroyl-D-quinic acid changing in R/FR (**Figure 8; Supp Figure 4-5**). Correspondingly, our metabolic pathway analysis, also revealed significant phenylpropanoid and flavonoid metabolic pathway enrichment associated with increasing R/B and R/FR (**Supp Table 3-4**). Here, we find a significant up-regulation of flavonoid biosynthetic pathways, which is more pronounced in the K9 cultivar and consistent with our ability to detect anthocyanins in this cultivar. Intriguingly, up-regulation of phenylpropanoid biosynthesis pathways in K9 with increasing R/B at ZT23, contrasts our observed anthocyanin levels, suggesting that at ZT23 the phenylpropanoid pathway may divert flux toward other phenylpropanoid products. For example, we see the coumarin biosynthesis pathway enriched in our metabolic pathway analysis (**Supp Table 3-4**), coupled with observed up-regulation of DEFECTIVE IN CUTICULAR RIDGES (DCR; A0A0D3DD85), ENHACED PSEUDOMONAS SUSCEPTIBILITY1 (EPS1; A0A0D3D884) and COUMARIN SYNTHASE (COSY; A0A0D3CEC8) in K9 at ZT23, indicating possible increased coumarin biosynthesis^56–58^. Thus, our results suggest the ideal conditions for kale anthocyanin production under the lower light intensities may involve use of a high R/FR spectral composition. This has substantial application potential for low energy indoor farming.

Although not measured in our metabolomics analysis, our proteomic analysis identified changes in abundance of proteins involved in production of glucosinolates and carotenoids (Figures 3-6). Individual glucosinolates are typically detected using HPLC, not GC-MS^59,60^ (**Figure 4-5**). Playing a role in kale flavour and plant defense, glucosinolates can also act as anti-oncogenic compounds for enhanced human health^61^. Our proteomic analyses indicate changes in glucosinolate metabolism with increasing light intensity and R/B ratio (**Figure 3**). In the K9 cultivar, increasing light intensity resulted in a down-regulation of proteins associated with glucosinolate catabolism. Our metabolic pathway analysis also suggests a potential for increased glucosinolate biosynthesis in K3 (**Supp Table 2**), thus the cultivars might achieve the same elevated levels of glucosinolates at higher light intensities through differing mechanisms. In our bioinformatics assessment of R/B responses in K3, we find decreased glucosinolate catabolism and increased glucosinolate biosynthesis (**Supp Table 3**). This has previously been observed in *B. napus* sprouts, where levels of the glucosinolates sinigrin, glucobrassicin, and 4-methoxy glucobrassicin were modulated by increasing red light levels^62^, while another study found differential cultivar responses in Chinese cabbage with changing light intensities^63^.

Lastly, carotenoids are also important antioxidant compounds beneficial to human health^64^. Within our proteomic data, we infer that K3 may have lower carotenoid production with increasing light intensity, as we quantified decreasing CAROTENE CIS-TRANS-ISOMERASE (CRTISO; A0A0D3DYP6), 15-CIS-PHYTOENE DESATURASE (PDS; A0A0D3BJ64) and PHYTOENE SYNTHASE1 (PSY1; A0A0D3AJG7) protein abundance. In our spectral composition experiments, it appears that increasing red light in both the R/B and R/FR experiments drives increased carotenoid biosynthesis in the K9 cultivar (**Figure 5**), as we see increasing CRTISO (A0A0D3DYP6), LYCOPENE EPSILON CYCLASE2 (LUT2; A0A0D3B320),

PSY1 and PDS protein abundance in R/B experiments and PDS, CRTISO and CYP97B3 protein abundance increases under R/FR conditions. These results suggest that K3 may be more sensitive to light intensity changes and K9 to levels of red light. We find it interesting to find a lower abundance of proteins involved in carotenoid biosynthesis with increasing light intensity in K3 as carotenoids tend to increase with higher light intensity^65^, however there is evidence in pepper of lower carotenoids with increasing light intensity that is suggested to be due to a higher photo-oxidation rate than that of carotenoid synthesis^66,67^. Structural diversity, degradation, volatility, and interactions with other compounds makes quantification of carotenoids difficult, and complex HPLC techniques are generally required^68^. Future studies looking to tailor the growth of kale for glucosinolate and/or carotenoid levels should look to quantify those metabolites directly, as that was beyond the scope of this study.

## Conclusions

Here, we present an integrated, systems-level analysis of two kale cultivars under increasing light intensity, R/B and R/FR spectral ratios with time-of-day precision. Our dataset reveals that light intensity and spectral composition can modulate myriad processes within kale, eliciting changes in growth and photosynthetic parameters, as well as the cell wall and hormones related to shade avoidance (**Figure 8**). Further, we identify changes in nutritional content such as amino acids, carbohydrates, glucosinolates, and anthocyanins that are of both fundamental and translational interest (**Figure 8**). Further investigation of signaling networks governing metabolism of interest would be of great interest. Collectively, these results provide new, wide-ranging knowledge to indoor / vertical farming initiatives of which genetics offer which nutritional benefits when grown under certain conditions and what time of day they may want to harvest them. This offers substantial value to indoor / vertical farming operations looking to grow diverse leafy greens of enhanced nutritional value.

## Materials and Methods

### Plant material and cultivation

Dwarf Curled Scotch (K3) and Red Scarlet (K9) kale seeds (*B. oleracea var. sabellica*) were purchased from West Coast Seeds (https://www.westcoastseeds.com/). Seeds were surface sterilized with incubations in 70% (v/v) ethanol and 30% (v/v) bleach solutions at room temperature beginning with a 4-minute incubation in 3 mL of 70% (v/v) ethanol. The seeds were then rinsed with 3 mL of sterile milli-Q water and 3 mL of 30% (v/v) bleach was added onto each tube for a 7-minute incubation with periodical shaking every 2 minutes. Finally, the bleach solution was decanted, and the seeds were rinsed three times with 3 mL of sterile Milli-Q water. Following sterilization, seeds from each cultivar were plated onto 0.5 x Murashige and Skoog (MS) media plates containing 1% (w/v) sucrose. The seeds were then stratified at 4 °C in total darkness for 4 days. Following stratification, the kale seedlings were entrained to their respective light treatment for 7 days. To prepare the soil for seedling transplant, ∼25 kg of soil (Sun Gro Sunshine Mix #1) was mixed with 8 L of H_2_O and then transferred onto 4” square polypropylene pots. Transplantation occurred once most plants had reached the cotyledon stage at 11 days post-imbibition (dpi).

### Light Conditions

The programmable LED (Perihelion Model) lights used for this experiment were provided by G2V Optics Inc. and were installed inside ventilated growth chambers lined with reflective mylar. The kale plants were grown under a 12-hour light and 12-hour dark regiment. Three different light intensities were tested: 75, 125, 175 PPFD (μmol.m^−2^.s^−1^) (**Figure 1A**). Spectral light conditions were assigned in spectral ratios of R/B and R/FR of low R/B or R/FR, balanced light and high R/B or R/FR (**Figure 1B**).

### Measurement of Kale Photosynthetic Performance via MultisynQ

Photosynthetic performance was measured following the Photosynthesis RIDES 2.0 protocol using a MultispeQ v1.0 (PhotosynQ LLC, USA)^30^. At mid-day (zeitgeber; ZT6) 34 dpi, the photosynthetic performance of the kale plants was measured by clamping the second true leaf between the MultisynQ sensors ensuring full coverage. The leaf remained in the chamber until the data collection process was completed (∼10 s). A total of 10 randomly selected plants were analyzed for each cultivar and light condition.

### Phenotyping measurements

Plants were grown in separate chambers under different light intensity and spectral conditions (**Figure 1A, 2A**). Each chamber was equipped with single-board Raspberry Pi 3 B+ computers and ArduCam Noir Cameras (OV5647 1080p) with a motorized IR-CUT filter and two infrared LEDs. Top-view pictures of plants were taken every 5 minutes. Plant surface area analysis across light conditions was performed by averaging the area of each plant measured during a 1-hour period (between ZT6 and ZT7) at 22 dpi. Plant surface areas were extracted using Python scripts from the open-source software package PlantCV (https://plantcv.readthedocs.io/), following the methodology previously described^69^. The pixel-based measurements were converted to cm in ImageJ by setting the known size of the plant pot as previously reported^70^.

### Harvesting and Sample Processing

At 35 dpi, kale leaves were harvested at two time points: end-of-night (ZT23) and end-of-day (ZT11). This was done to capture the expansive metabolic changes occurring during the transitions between night-to-day and day-to-night, respectively. All leaves from three random kale plants of the same cultivar were pooled into a labelled 50 mL conical tube and promptly frozen in liquid nitrogen to preserve the proteins and metabolites present. In total, four biological reps (n=4) were collected from each cultivar per chamber during harvest. Two metal beads were then introduced into each conical tube and the samples were ground by subjecting them to 3x – 30 s/1200 rpm using a Geno/Grinder^®^ - Automated Tissue Homogenizer and Cell Lyser. 100 mg (±1 mg) of each finely-ground sample was then portioned into each of two separate 2 mL Eppendorf Safe-Lock tubes for further processing (one for proteomics, the other for metabolomics).

### Anthocyanin Extraction and Quantification

Anthocyanin analysis was performed as previously described by Neff and Chory (1998), with the following modifications. Specifically, 500 µL of 100% (v/v) methanol-1% (v/v) hydrochloric acid was added to each tube containing the finely ground samples and incubated at 4 °C for 24 hours in darkness. From there, 200 µL of H_2_O and 500 µL of chloroform were added and the tubes were centrifuged at 18,000 x g for 5 minutes at room temperature. 400 µL of the supernatant was then transferred over to new tubes and adjusted to a total volume of 800 µL with 60% (v/v) methanol-1% (v/v) hydrochloric acid. A total of 200 µL from each volume-corrected sample was transferred into a well in a 96-well plate and absorbance at 530 nm and 657 nm was measured with a Tecan Spark® plate reader using 60% (v/v) methanol −1% (v/v) hydrochloric acid as the background. The relative anthocyanin content was then derived mathematically using: Antho = (Abs_530_ - Abs_657_) x 1000 x powder weight (mg)^-1^.

### Metabolite analysis via Gas Chromatography Mass Spectrometry (GCMS)

*Metabolite extraction:* A total of 700 µL of ice-cold 100% (v/v) methanol was added to the tubes containing 100 mg of finely ground leaf sample and promptly vortexed for ∼20 seconds. Using a thermomixer, these were then incubated for 15 minutes at 70 °C with shaking at 1200 rpm. From there, the tubes were centrifuged at 18,000 rpm for 15 minutes and the supernatants were transferred into new tubes. The pellet was then re-extracted using 700 µL of 50% (v/v) methanol containing Ribitol at 25 µL per sample at 0.4 mg/mL in water and vortexed for ∼20 seconds until re-suspended. These tubes were then centrifuged at 14,000 rpm for 10 minutes and 600 µL of the supernatant was combined with the previous supernatant. Finally, 100 µL was transferred into new tubes and dried using a vacuum centrifuge at 1 torr at room temperature for 1.5 - 2 hours.

### Gas chromatography mass spectrometry analysis

The dried samples were derivatized by adding 100 µL of methoxamine hydrochloride into each tube and then incubated at 30 °C at 850 rpm for 1 hour using a Thermomixer F2.0 (Eppendorf). Subsequently, 50 µL of N,O-bis(trimethylsilyl)trifluoroacetamide (BSTFA) was added to each tube and incubated at 50 °C at 850 rpm for 3 hours using a Thermomixer F2.0 (Eppendorf). A total of 150 µL of each sample was then transferred to vials in preparation for injection in the GC-MS. Samples were analyzed via a 7890A gas chromatograph (Agilent) paired to a 5975C quadrupole mass detector (Aglient). The initial GC oven temperature was set at 70 °C and then increased to 325 °C at 7 °C per minute two minutes post-injection where the temperature was then maintained at 325 °C for 3.6 minutes. The injection and ion source temperatures were set to 300 °C and 230 °C, respectively with a solvent delay of 5 minutes. The helium flow rate was also adjusted to a rate of 1 mL/minute. Measurements were performed with electron impact ionization (70 eV) in full scan mode (m/z 33-600). Identification of metabolites was performed based on their mass spectral and retention time index, and how closely they matched to those acquired from the National Institute of Standards and Technology library and the Golm Metabolome Database.

### Protein extraction and LC-MS analysis

Kale leaf tissue harvested at ZT23 and ZT11 was flash-frozen in liquid N_2_. Tissue was ground in liquid N_2_ using a mortar and pestle and aliquoted into 400 mg samples (n=4 replicates). Samples were extracted with 50 mM HEPES-KOH pH 8.0, 50 mM NaCl, and 4% (w/v) SDS at a ratio of 1:2 (w/v). Next, samples were vortexed followed by incubation for 15 min at 95 °C using a Thermomixer F2.0 (Eppendorf) shaking at 1,100 rpm. Samples were subsequently incubated at room temperature for an additional 15 min of shaking followed by clarification at 20,000 x g for 5 min at room temperature, where the supernatant was then retained in fresh Eppendorf microtubes. Protein concentration was estimated by bicinchoninic (BCA) assay (23225; Thermo Scientific), followed by sample reduction using 10 mM dithiothreitol (DTT) for 5 min at 95 °C. Following cooling to room temperature, samples were alkylated using 30 mM iodoacetamide (IA) in dark for 30 min at room temperature without shaking. The alkylation was stopped by addition of 10 mM DTT, followed by a brief vortex and 10 min room temperature incubation without shaking. Samples were digested overnight with 1:100 sequencing grade trypsin (V5113; Promega), followed by quantification of the generated peptide pools using a Nanodrop (Thermo Scientific). Subsequently, samples were acidified with formic acid to a final concentration of 5% (v/v) followed by drying by vacuum centrifugation. Peptide desalting was then performed using ZipTip C18 pipette tips (ZTC18S960; Millipore), with peptides subsequently dried and dissolved in 3% (v/v) ACN / 0.1% (v/v) FA prior to MS analysis. A Fusion Lumos Tribrid Orbitrap mass spectrometer (Thermo Scientific) was used to analyse the digested peptides in data independent acquisition (DIA) mode using the BoxCarDIA method as previously described^71^. 1 µg of dissolved peptide was injected using an Easy-nLC 1200 system (LC140; ThermoScientific) and a 50 cm Easy-Spray PepMap C18 column (ES903; ThermoScientific). Liquid chromatography and BoxCarDIA acquisition was performed as previously described^71^.

### Proteomic Data Analysis

All acquired BoxCarDIA data was analyzed using a library-free approach in Spectronaut v14 (Biognosys AG) under default settings. The *B. oleracea* var. *oleracea* proteome (Uniprot:https://www.uniprot.org/ containing 58,545 proteins) was used for data searching. Default search parameters were used for proteome quantification including: a protein, peptide and PSM FDR of 1%, trypsin digestion with 1 missed cleavage, fixed modification including carbamidomethylation of cysteine residues and variable modification including methionine oxidation. Data was Log2 transformed and median normalized with significantly changing differentially abundant proteins determined and corrected for multiple comparisons (Bonferroni-corrected p-value <0.05; q-value). Each biological replicate consisted of pools of leaves from 3 plants, with each condition consisting of n=4 biological replicates.

### Bioinformatics

Gene Ontology (GO) enrichment analysis was performed using the Database for Annotation, Visualization and Integrated Discovery (DAVID; v 6.8; https://david.ncifcrf.gov/home.jsp)). A using a significance threshold of p-value < 0.01. Conversion of *B. oleracea* gene identifiers to Arabidopsis was performed using Uniprot (https://www.uniprot.org/) followed by Ensembl Biomart (https://plants.ensembl.org/biomart) for STRING association network analysis and metabolic pathway analysis. STRING association network analysis was performed using Cytoscape v3.10.1 (https://cytoscape.org) in combination with the String DB plugin stringApp and an edge threshold of 0.9 for Intensity and 0.8 for Spectra R/B and R/FR. Visualization of GO dotplot was performed using R version 4.3.1 and *ggplot2* package. Metabolic pathway enrichment was performed using the Plant Metabolic Network (PMN; https://plantcyc.org) with enrichment determined using a Fisher’s exact test (p-value < 0.01). Final figures were assembled using Affinity Designer (v.2.4.1).

### Phenotyping statistical analysis

Morphological statistical analysis was done as stated in the figure captions. For the most part, these are based on a one-way ANOVA followed by a Tukey *post-hoc* test (corrected p-value ≤ 0.05).

## DATA AVAILABILITY

All raw data files have been uploaded to ProteomeXchange (http://www.proteomexchange.org/) via the PRoteomics IDEntification Database (PRIDE; https://www.ebi.ac.uk/pride/) with the dataset identifier PXD055752.

## ACKNOWLEDGEMENTS

The authors thank the Natural Sciences and Engineering Research Council of Canada (NSERC), Alberta Innovates (AI) and Canada Foundation for Innovation (CFI) for funding. The authors thank G2V Optics Inc. for providing their programmable LED lighting system as well as Jack Moore of the Alberta Proteomics and Mass Spectrometry Facility for assistance with mass spectrometer operation and maintenance.

## AUTHOR CONTRIBUTIONS

R.G.U. and S.S. conceptualized the study. S.S., B.C., L.I., C.K., N.B., M.T. performed research and data acquisition. S.S., L.E.G. and R.G.U. analyzed the data and wrote the paper with input from all authors.

---

**Supplemental Figure 1.**
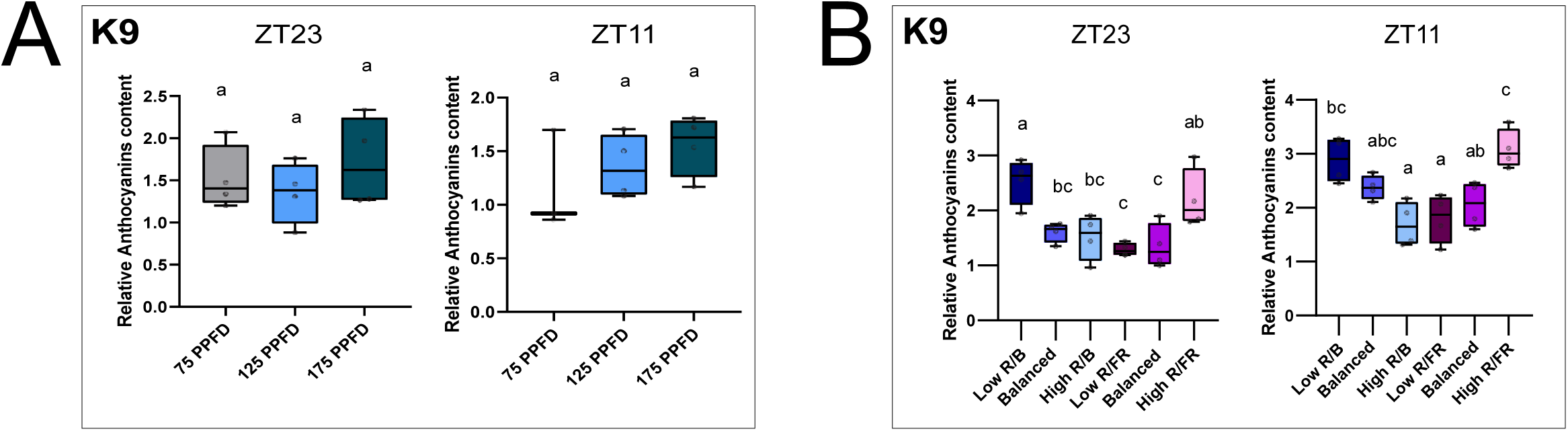
Effect of light intensity and spectra on K9 anthocyanin content. A. K9 anthocyanin content at 35 dpi (n=4), at different light intensities (75, 125, and 175 PPFD). B. K9 anthocyanin content at 35 dpi (n=4), at different light spectra (low, balanced, high R/B and R/FR). Letters shows significant differences using a one-way ANOVA and Tukey’s post-hoc test (adjusted p-value <0.05).

**Supplemental Figure 2.**
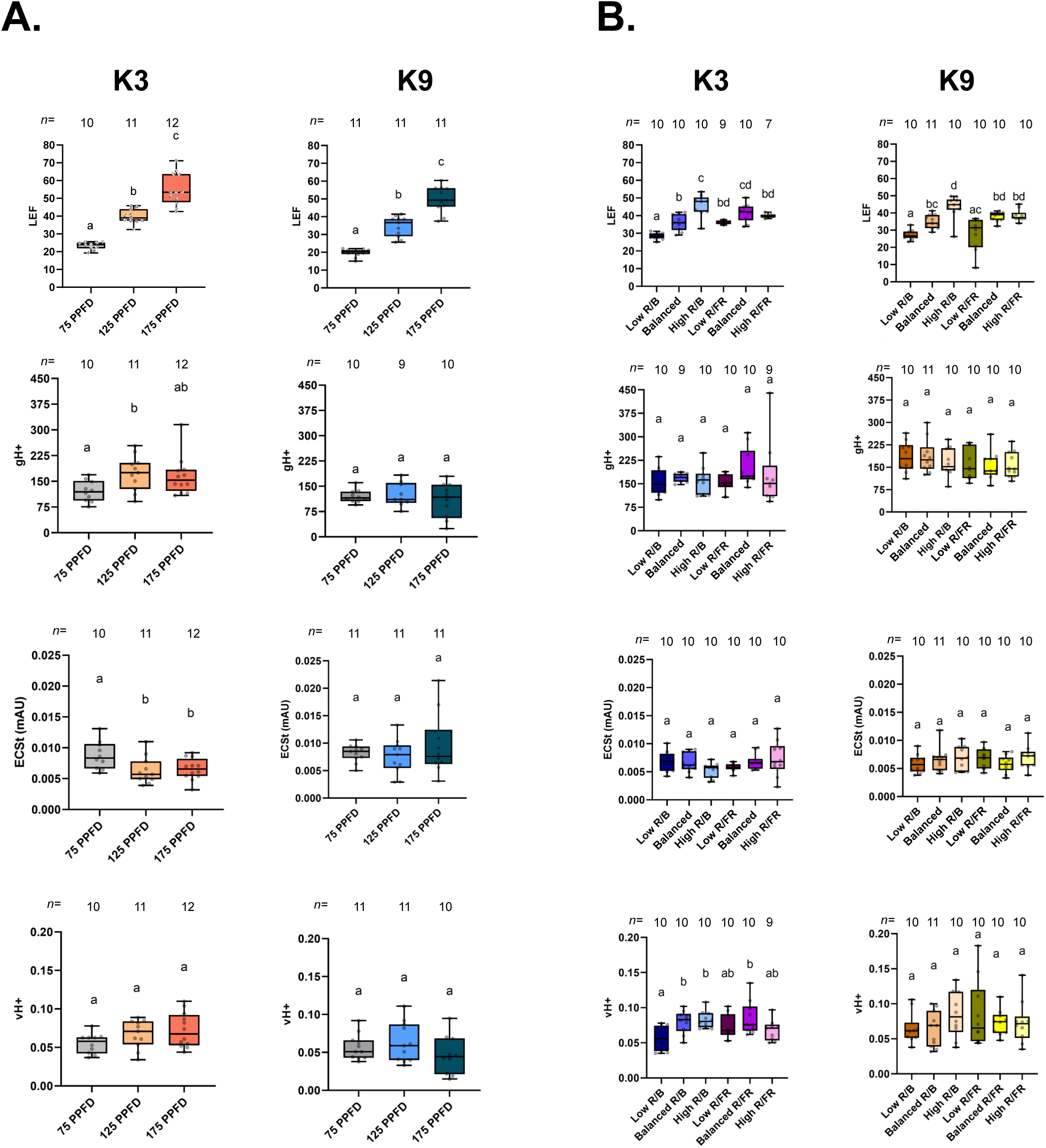
Effect of Light Intensity and spectra on Photosynthetic Parameters Measured with MultispeQ. A. Light intensity effect on photosynthetic parameters (LEF, gH+, ECSt, vH+) were measured with MultispeQ at 35 dpi (n=10). B. Light spectra effect on photosynthetic parameters (LEF, gH+, ECSt, vH+) were measured with MultispeQ at 35 dpi (n=10). Letters show significant differences using a one-way ANOVA and Tukey’s post-hoc test (adjusted p-value < 0.05).

**Supplemental Figure 3.**
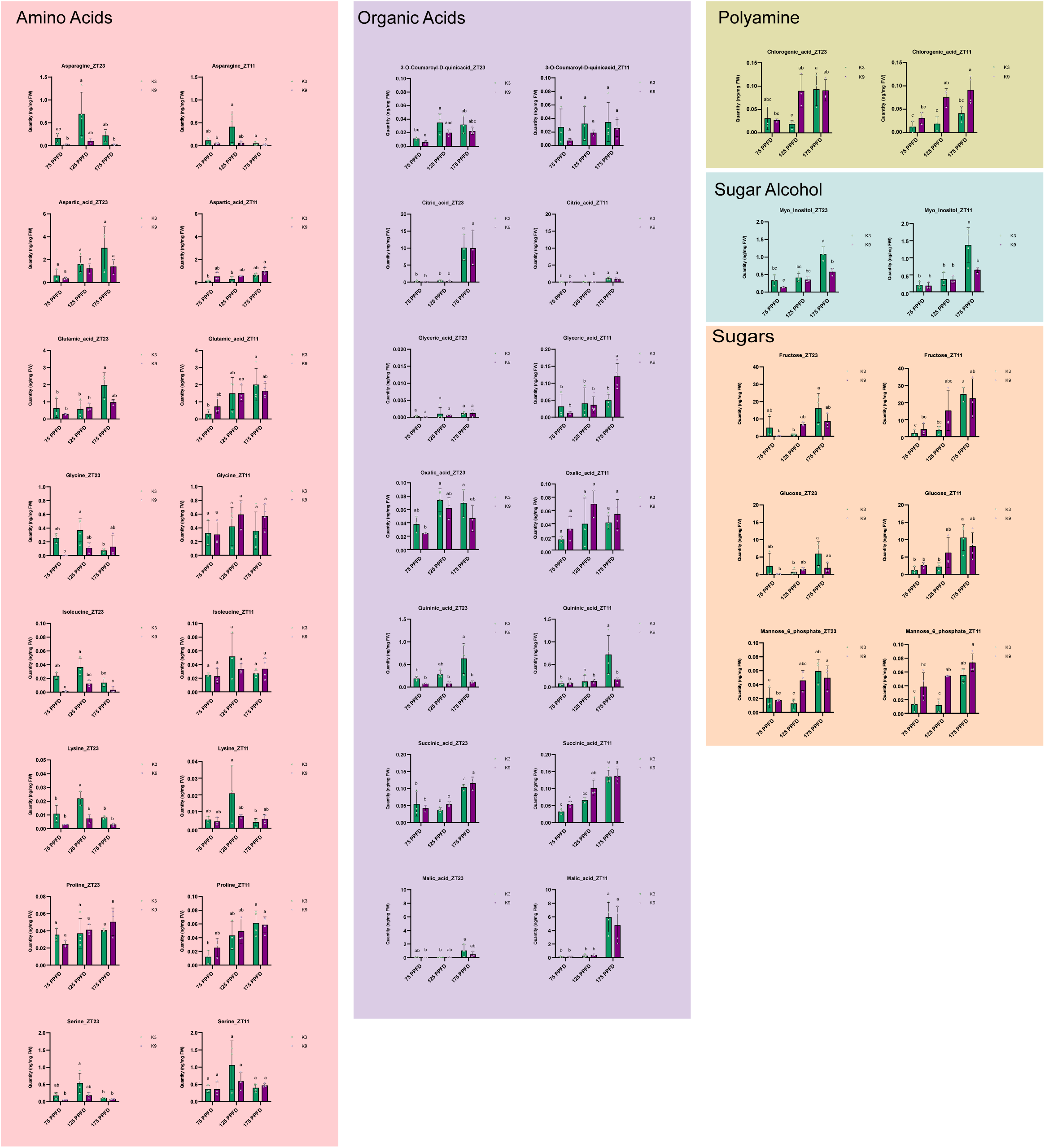
Effect of light intensity treatment on the metabolite composition of Kale cultivars. GC-MS analysis of primary metabolites in kale cultivars K3 and K9 under different light intensity 75, 125 and 175 PPFD and at ZT11 / ZT23 (n=4). Different letters indicate significant differences using a one-way ANOVA and Tukey’s post-hoc test (adjusted p-value < 0.05).

**Supplemental Figure 4.**
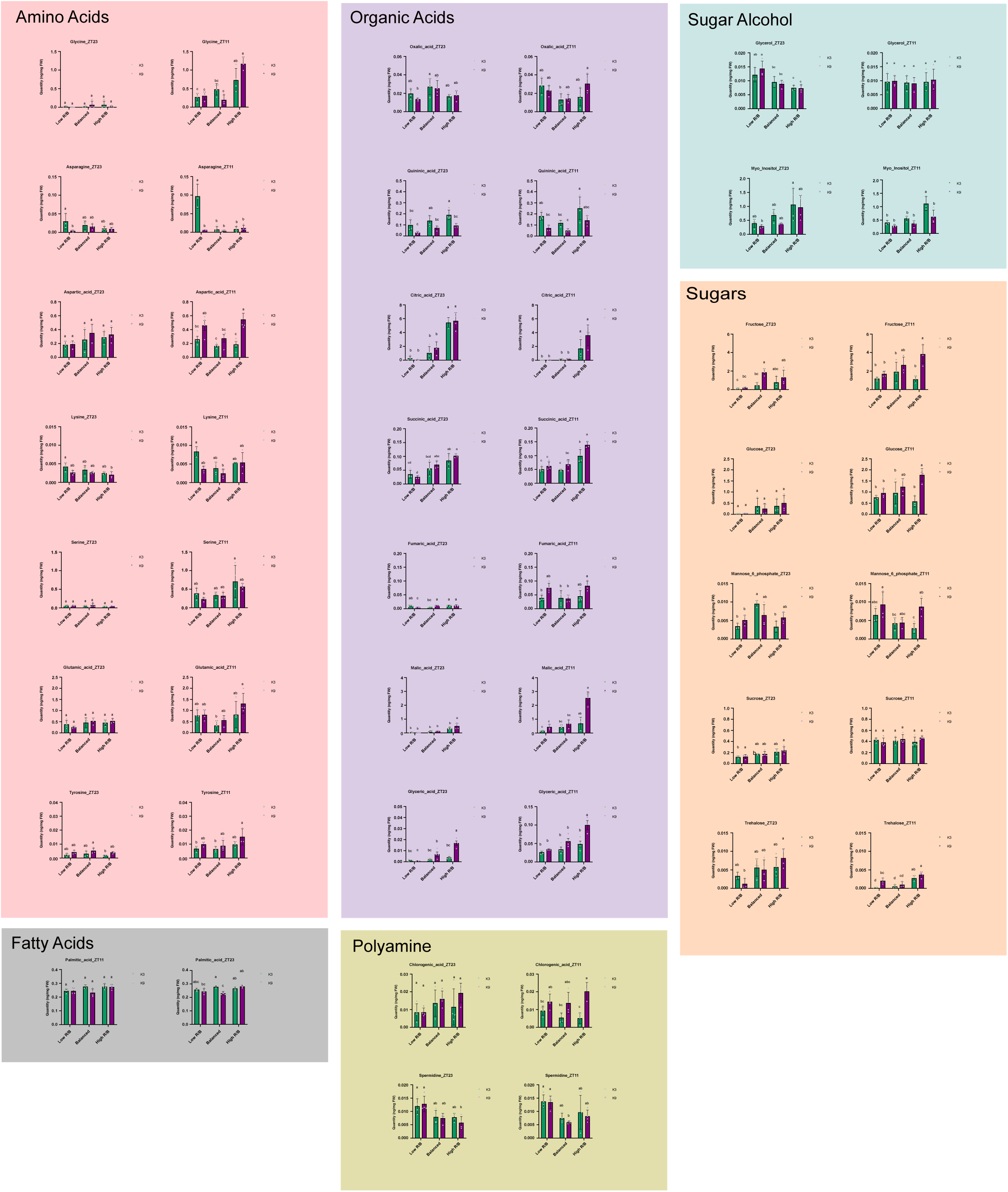
Effect of light spectra treatment on the metabolite composition of Kale cultivars. GC-MS analysis of primary metabolites in kale cultivars K3 and K9 under different light spectra (low, balanced and high R/B) and at ZT11 / ZT23 (n=4). Different letters indicate significant differences using a one-way ANOVA and Tukey’s post-hoc test (adjusted p-value < 0.05).

**Supplemental Figure 5.**
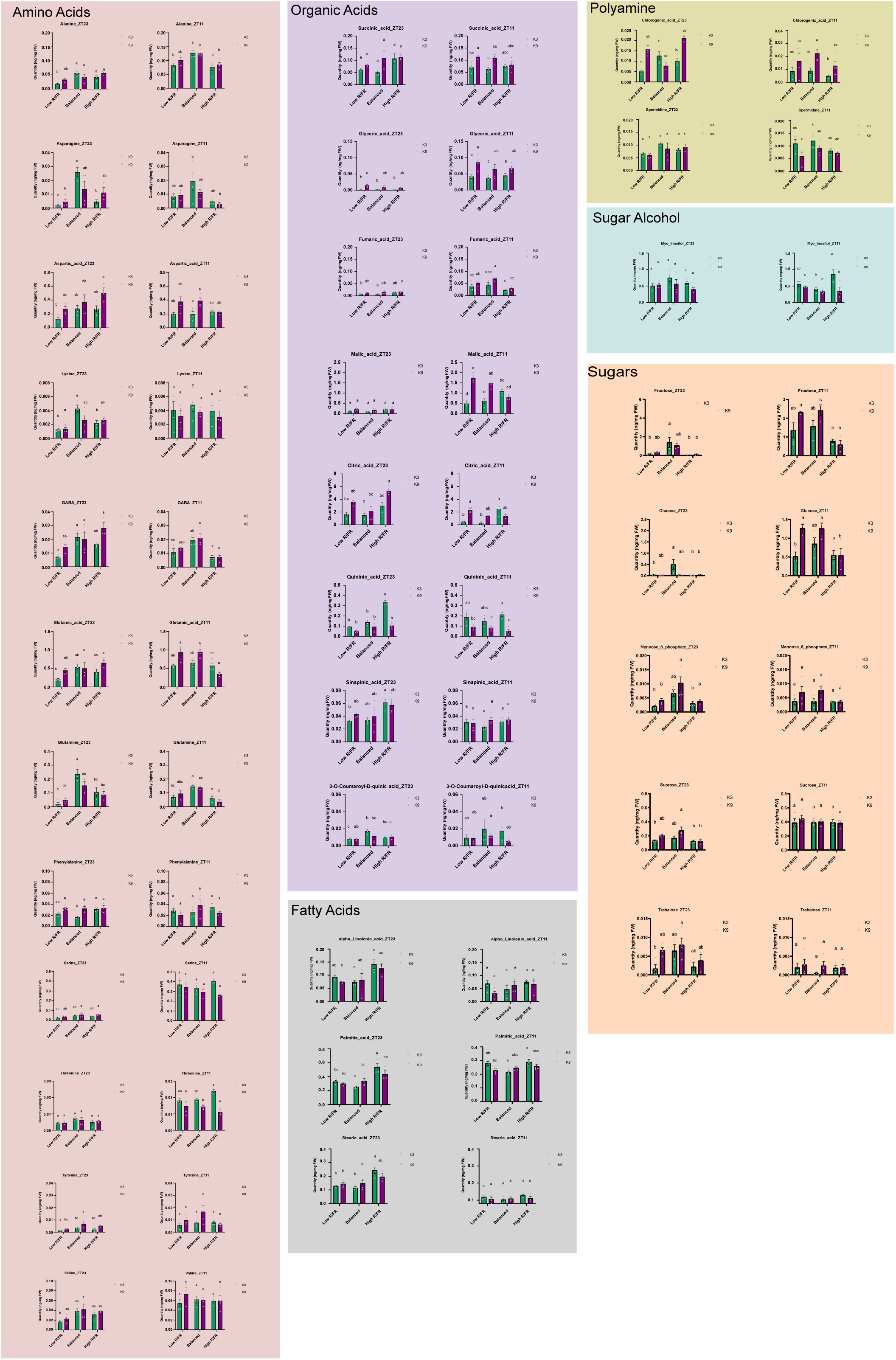
Effect of light spectra treatment on the metabolite composition of Kale cultivars. GC-MS analysis of primary metabolites in kale cultivars K3 and K9 under different light spectra (low, balanced and high R/FR) and at ZT11 / ZT23 (n=4). Different letters indicate significant differences using a one-way ANOVA and Tukey’s post-hoc test (adjusted p-value < 0.05).

**Supplemental Figure 6.**
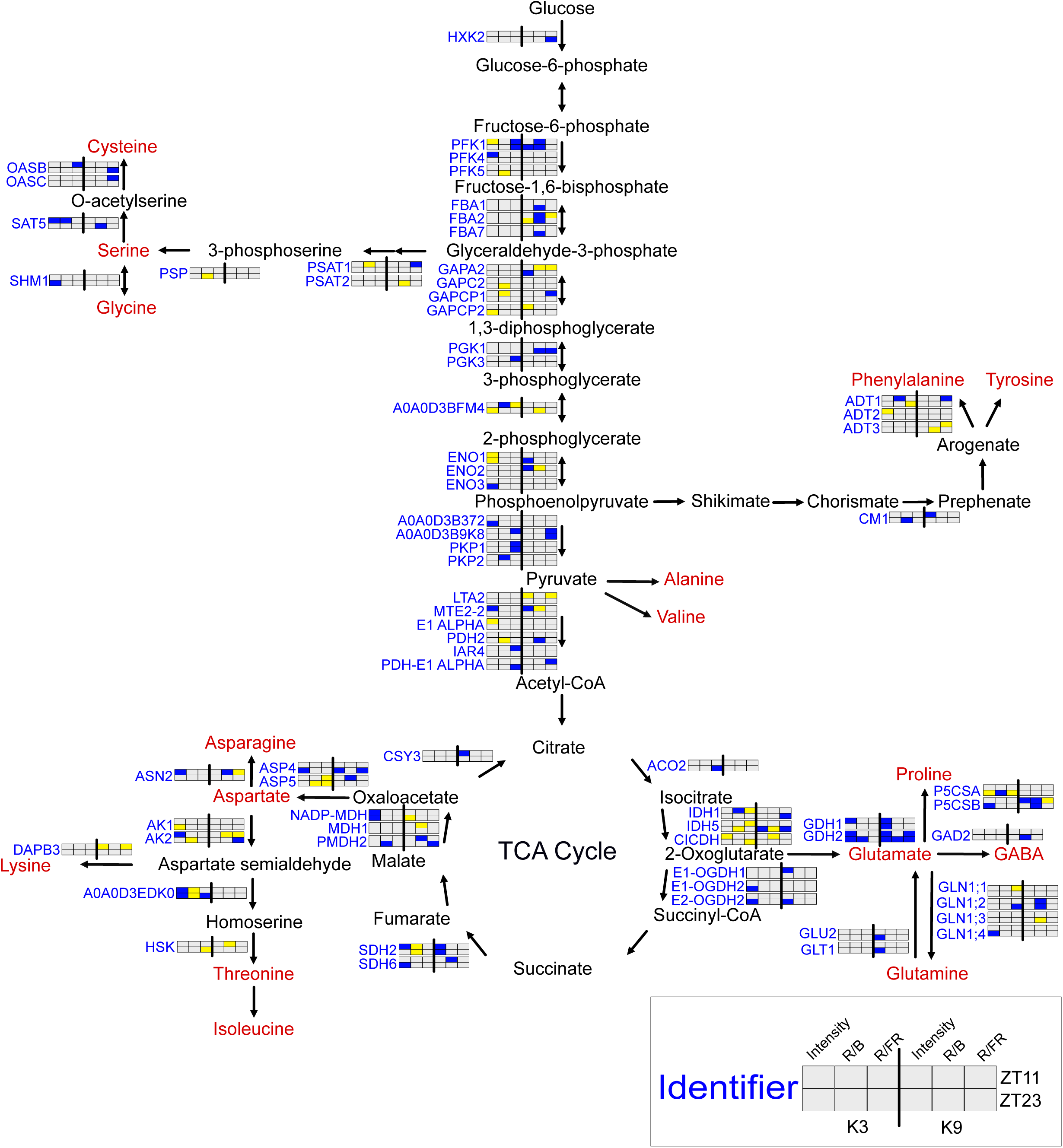
Summary of proteome changes related to amino acid metabolism. Schematic depicting amino acid metabolism related proteome changes in K3 and K9 at ZT11 and ZT23. Blue lettering indicates proteins that are significantly changing. Blue and yellow boxes indicate down- and up-regulation, respectively. Red lettering denotes amino acids that were measured in metabolite analysis.

**Supplemental Table 1.** All quantified and significantly changing proteins with changing light intensity and spectral composition

**Supplemental Table 2.** Pathway enrichment analysis using the Plant Metabolic Network (PMN; https://plantcyc.org) for Intensity experiments

**Supplemental Table 3.** Pathway enrichment analysis using the Plant Metabolic Network (PMN; https://plantcyc.org) for R/B Spectra experiments

**Supplemental Table 4.** Pathway enrichment analysis using the Plant Metabolic Network (PMN; https://plantcyc.org) for R/FR Spectra experiments

**Supplemental Table 5.** Gas chromatography metabolite data for Intensity experiments

**Supplemental Table 6.** Gas chromatography metabolite data for Spectra experiments

## REFERENCES

1. Satheesh, N. & Workneh Fanta, S. Kale: Review on nutritional composition, bio-active compounds, anti-nutritional factors, health beneficial properties and value-added products. Cogent Food Agric. 6, 1811048 (2020).

2. Rachwał, K. et al. Red Kale (Brassica oleracea L. ssp. acephala L. var. sabellica) Induces Apoptosis in Human Colorectal Cancer Cells In Vitro. Molecules 28, 6938 (2023).

3. Bowen-Forbes, C. et al. Broccoli, Kale, and Radish Sprouts: Key Phytochemical Constituents and DPPH Free Radical Scavenging Activity. Molecules 28, 4266 (2023).

4. Cao, D. et al. Spermidine enhances chilling tolerance of kale seeds by modulating ROS and phytohormone metabolism. PLOS ONE 18, e0289563 (2023).

5. Bauer, N. et al. Mechanisms of Kale (Brassica oleracea var. acephala) Tolerance to Individual and Combined Stresses of Drought and Elevated Temperature. Int. J. Mol. Sci. 23, 11494 (2022).

6. Fussy, A. & Papenbrock, J. An Overview of Soil and Soilless Cultivation Techniques—Chances, Challenges and the Neglected Question of Sustainability. Plants 11, 1153 (2022).

7. https://www.alliedmarketresearch.com, A. M. R. Kale Microgreen Market Size, Share, Price, Trends |Research Report 2030. *Allied Market Research* https://www.alliedmarketresearch.com/kale-microgreen-market-A16137.

8. Hati, A. J. & Singh, R. R. Smart Indoor Farms: Leveraging Technological Advancements to Power a Sustainable Agricultural Revolution. AgriEngineering 3, 728–767 (2021).

9. Nájera, C., Gallegos-Cedillo, V. M., Ros, M. & Pascual, J. A. LED Lighting in Vertical Farming Systems Enhances Bioactive Compounds and Productivity of Vegetables Crops. Biol. Life Sci. Forum 16, 24 (2022).

10. McCree, K. J. The action spectrum, absorptance and quantum yield of photosynthesis in crop plants. Agric. Meteorol. 9, 191–216 (1971).

11. Krahmer, J., Ganpudi, A., Abbas, A., Romanowski, A. & Halliday, K. J. Phytochrome, Carbon Sensing, Metabolism, and Plant Growth Plasticity. Plant Physiol. 176, 1039–1048 (2018).

12. Chen, J. et al. Effect of Photoperiod on Chinese Kale (Brassica alboglabra) Sprouts Under White or Combined Red and Blue Light. Front. Plant Sci. 11, (2021).

13. Kong, Y., Schiestel, K. & Zheng, Y. Pure blue light effects on growth and morphology are slightly changed by adding low-level UVA or far-red light: A comparison with red light in four microgreen species. Environ. Exp. Bot. 157, 58–68 (2019).

14. Deng, M. et al. Influence of pre-harvest red light irradiation on main phytochemicals and antioxidant activity of Chinese kale sprouts. Food Chem. 222, 1–5 (2017).

15. Zhen, S. & van Iersel, M. W. Far-red light is needed for efficient photochemistry and photosynthesis. J. Plant Physiol. 209, 115–122 (2017).

16. Furuya, M. & Schäfer, E. Photoperception and signalling of induction reactions by different phytochromes. Trends Plant Sci. 1, 301–307 (1996).

17. Scandola, S., Mehta, D., Castillo, B., Boyce, N. & Uhrig, R. G. Systems-level proteomics and metabolomics reveals the diel molecular landscape of diverse kale cultivars. Front. Plant Sci. 14, (2023).

18. Boucher, L., Nguyen, T.-T.-A., Brégard, A., Pepin, S. & Dorais, M. Optimizing Light Use Efficiency and Quality of Indoor Organically Grown Leafy Greens by Using Different Lighting Strategies. Agronomy 13, 2582 (2023).

19. Balázs, L. et al. Quantifying the Effect of Light Intensity Uniformity on the Crop Yield by Pea Microgreens Growth Experiments. Horticulturae 9, 1187 (2023).

20. Jones-Baumgardt, C., Llewellyn, D., Ying, Q. & Zheng, Y. Intensity of Sole-source Light-emitting Diodes Affects Growth, Yield, and Quality of Brassicaceae Microgreens. 10.21273/HORTSCI13788-18 (2019) doi:10.21273/HORTSCI13788-18.

21. Dou, H. et al. Supplementary Far-Red Light for Photosynthetic Active Radiation Differentially Influences the Photochemical Efficiency and Biomass Accumulation in Greenhouse-Grown Lettuce. Plants 13, 2169 (2024).

22. Gudžinskaitė, I., Laužikė, K., Pukalskas, A. & Samuolienė, G. The Effect of Light Intensity during Cultivation and Postharvest Storage on Mustard and Kale Microgreen Quality. Antioxidants 13, 1075 (2024).

23. Boucher, L., Nguyen, T.-T.-A., Brégard, A., Pepin, S. & Dorais, M. Optimizing Light Use Efficiency and Quality of Indoor Organically Grown Leafy Greens by Using Different Lighting Strategies. Agronomy 13, 2582 (2023).

24. Zhang, Y., Butelli, E. & Martin, C. Engineering anthocyanin biosynthesis in plants. Curr. Opin. Plant Biol. 19, 81–90 (2014).

25. Waterland, N. L. et al. Differences in Leaf Color and Stage of Development at Harvest Influenced Phytochemical Content in Three Cultivars of Kale (Brassica oleracea L. and B. napus). J. Agric. Sci. 11, p14 (2019).

26. Kaur, S. et al. Protective and defensive role of anthocyanins under plant abiotic and biotic stresses: An emerging application in sustainable agriculture. J. Biotechnol. 361, 12–29 (2023).

27. Dou, H., Niu, G. & Gu, M. Photosynthesis, Morphology, Yield, nd Phytochemical Accumulation in Basil Plants Influenced by Substituting Green Light for Partial Red and/or Blue Light. 10.21273/HORTSCI14282-19 (2019) doi:10.21273/HORTSCI14282-19.

28. Anum, H., Cheng, R. & Tong, Y. Improving plant growth, anthocyanin production and oxidative status of red lettuce (*Lactuca sativa* cv. Lolla Rossa) by optimizing red to blue light ratio with a constant green light fraction in a plant factory. Sci. Hortic. 338, 113832 (2024).

29. Frąszczak, B. et al. Morphological and Photosynthetic Parameters of Green and Red Kale Microgreens Cultivated under Different Light Spectra. Plants 12, 3800 (2023).

30. Kuhlgert, S. et al. MultispeQ Beta: a tool for large-scale plant phenotyping connected to the open PhotosynQ network. R. Soc. Open Sci. 3, 160592 (2016).

31. Fu, W., Li, P. & Wu, Y. Effects of different light intensities on chlorophyll fluorescence characteristics and yield in lettuce. Sci. Hortic. 135, 45–51 (2012).

32. Miao, Y., Wang, X., Gao, L., Chen, Q. & Qu, M. Blue light is more essential than red light for maintaining the activities of photosystem II and I and photosynthetic electron transport capacity in cucumber leaves. J. Integr. Agric. 15, 87–100 (2016).

33. Pierik, R. & de Wit, M. Shade avoidance: phytochrome signalling and other aboveground neighbour detection cues. J. Exp. Bot. 65, 2815–2824 (2014).

34. Lv, M. & Li, J. Molecular Mechanisms of Brassinosteroid-Mediated Responses to Changing Environments in Arabidopsis. Int. J. Mol. Sci. 21, 2737 (2020).

35. Keuskamp, D. H. et al. Blue-light-mediated shade avoidance requires combined auxin and brassinosteroid action in Arabidopsis seedlings. Plant J. 67, 208–217 (2011).

36. Klahre, U. et al. The Arabidopsis DIMINUTO/DWARF1 Gene Encodes a Protein Involved in Steroid Synthesis. Plant Cell 10, 1677–1690 (1998).

37. Ma, L. & Li, G. Auxin-Dependent Cell Elongation During the Shade Avoidance Response. Front. Plant Sci. 10, (2019).

38. Sasidharan, R., Chinnappa, C. C., Voesenek, L. A. C. J. & Pierik, R. The Regulation of Cell Wall Extensibility during Shade Avoidance: A Study Using Two Contrasting Ecotypes of Stellaria longipes. Plant Physiol. 148, 1557–1569 (2008).

39. Lisiewska, Z., Kmiecik, W. & Korus, A. The amino acid composition of kale (*Brassica oleracea* L. var. *acephala*), fresh and after culinary and technological processing. Food Chem. 108, 642–648 (2008).

40. Okumoto, S., Funck, D., Trovato, M. & Forlani, G. Editorial: Amino Acids of the Glutamate Family: Functions beyond Primary Metabolism. Front. Plant Sci. 7, (2016).

41. Kavi Kishor, P. B. & Sreenivasulu, N. Is proline accumulation per se correlated with stress tolerance or is proline homeostasis a more critical issue? Plant Cell Environ. 37, 300–311 (2014).

42. Galili, G. & Amir, R. Fortifying plants with the essential amino acids lysine and methionine to improve nutritional quality. Plant Biotechnol. J. 11, 211–222 (2013).

43. Yang, Q., Zhao, D. & Liu, Q. Connections Between Amino Acid Metabolisms in Plants: Lysine as an Example. Front. Plant Sci. 11, (2020).

44. Hall, C. J. et al. Differential lysine-mediated allosteric regulation of plant dihydrodipicolinate synthase isoforms. FEBS J. 288, 4973–4986 (2021).

45. Van Bochaute, P., Novoa, A., Ballet, S., Rognes, S. E. & Angenon, G. Regulatory mechanisms after short- and long-term perturbed lysine biosynthesis in the aspartate pathway: the need for isogenes in Arabidopsis thaliana. Physiol. Plant. 149, 449–460 (2013).

46. Ma, Q., Cao, X., Wu, L., Mi, W. & Feng, Y. Light intensity affects the uptake and metabolism of glycine by pakchoi (Brassica chinensis L.). Sci. Rep. 6, 21200 (2016).

47. Rosa-Téllez, S. et al. The serine–glycine–one-carbon metabolic network orchestrates changes in nitrogen and sulfur metabolism and shapes plant development. Plant Cell 36, 404–426 (2024).

48. Hwang, J. E. et al. Distinct expression patterns of two Arabidopsis phytocystatin genes, AtCYS1 and AtCYS2, during development and abiotic stresses. Plant Cell Rep. 29, 905–915 (2010).

49. Avidan, O. et al. Direct and indirect responses of the Arabidopsis transcriptome to an induced increase in trehalose 6-phosphate. Plant Physiol. 196, 409–431 (2024).

50. Sharma, N., Chaudhary, C. & Khurana, P. Role of myo-inositol during skotomorphogenesis in Arabidopsis. Sci. Rep. 10, 17329 (2020).

51. Gęgotek, A. & Skrzydlewska, E. Antioxidative and Anti-Inflammatory Activity of Ascorbic Acid. Antioxidants 11, 1993 (2022).

52. Pashkovskiy, P. et al. Post-Harvest Red- and Far-Red-Light Irradiation and Low Temperature Induce the Accumulation of Carotenoids, Capsaicinoids, and Ascorbic Acid in Capsicum annuum L. Green Pepper Fruit. Foods 12, 1715 (2023).

53. Tena, N., Martín, J. & Asuero, A. G. State of the Art of Anthocyanins: Antioxidant Activity, Sources, Bioavailability, and Therapeutic Effect in Human Health. Antioxidants 9, 451 (2020).

54. Pojer, E., Mattivi, F., Johnson, D. & Stockley, C. S. The Case for Anthocyanin Consumption to Promote Human Health: A Review. Compr. Rev. Food Sci. Food Saf. 12, 483–508 (2013).

55. Liu, C., Yao, X., Li, G., Huang, L. & Xie, Z. Transcriptomic profiling of purple broccoli reveals light-induced anthocyanin biosynthetic signaling and structural genes. PeerJ 8, e8870 (2020).

56. Kim, C. Y. et al. Emergence of a proton exchange-based isomerization and lactonization mechanism in the plant coumarin synthase COSY. Nat. Commun. 14, 597 (2023).

57. Vanholme, R. et al. COSY catalyses trans–cis isomerization and lactonization in the biosynthesis of coumarins. Nat. Plants 5, 1066–1075 (2019).

58. Rani, S. H., Krishna, T. H. A., Saha, S., Negi, A. S. & Rajasekharan, R. Defective in Cuticular Ridges (*DCR*) of *Arabidopsis thaliana*, a Gene Associated with Surface Cutin Formation, Encodes a Soluble Diacylglycerol Acyltransferase*. J. Biol. Chem. 285, 38337–38347 (2010).

59. Grosser, K. & van Dam, N. M. A Straightforward Method for Glucosinolate Extraction and Analysis with High-pressure Liquid Chromatography (HPLC). J. Vis. Exp. JoVE 55425 (2017) doi:10.3791/55425.

60. Theunis, M. et al. Optimization and validation of analytical RP-HPLC methods for the quantification of glucosinolates and isothiocyanates in *Nasturtium officinale* R. Br and Brassica oleracea. LWT 165, 113668 (2022).

61. Ltd, wflpublisher. Comparative study on functional components, antioxidant activity and color parameters of selected colored leafy vegetables as affected by photoperiods. wflpublisher.com https://www.wflpublisher.com/Abstract/2605.

62. Park, C. H. et al. Effects of Light-Emitting Diodes on the Accumulation of Glucosinolates and Phenolic Compounds in Sprouting Canola (Brassica napus L.). Foods 8, 76 (2019).

63. Zhou, B. et al. Effects of light intensity on the biosynthesis of glucosinolate in Chinese cabbage plantlets. Sci. Hortic. 316, 112036 (2023).

64. Eggersdorfer, M. & Wyss, A. Carotenoids in human nutrition and health. Arch. Biochem. Biophys. 652, 18–26 (2018).

65. Flores, M., Hernández-Adasme, C., Guevara, M. J. & Escalona, V. H. Effect of different light intensities on agronomic characteristics and antioxidant compounds of Brassicaceae microgreens in a vertical farm system. Front. Sustain. Food Syst. 8, (2024).

66. Simkin, A. J., Zhu, C., Kuntz, M. & Sandmann, G. Light-dark regulation of carotenoid biosynthesis in pepper (*Capsicum annuum*) leaves. J. Plant Physiol. 160, 439–443 (2003).

67. Pizarro, L. & Stange, C. Light-dependent regulation of carotenoid biosynthesis in plants. Cienc. E Investig. Agrar. 36, 143–162 (2009).

68. Kurek, M. A., Aktaş, H., Pokorski, P., Pogorzelska-Nowicka, E. & Custodio-Mendoza, J. A. A Comprehensive Review of Analytical Approaches for Carotenoids Assessment in Plant-Based Foods: Advances, Applications, and Future Directions. Appl. Sci. 15, 3506 (2025).

69. Gehan, M. A. et al. PlantCV v2: Image analysis software for high-throughput plant phenotyping. PeerJ 5, e4088 (2017).

70. Sakeef, N. et al. Machine learning classification of plant genotypes grown under different light conditions through the integration of multi-scale time-series data. Comput. Struct. Biotechnol. J. 21, 3183–3195 (2023).

71. Mehta, D., Scandola, S. & Uhrig, R. G. BoxCar and Library-Free Data-Independent Acquisition Substantially Improve the Depth, Range, and Completeness of Label-Free Quantitative Proteomics. Anal. Chem. 94, 793–802 (2022).

